# Molecular dynamics simulations and *in vivo* binding studies indicate different binding modes of plant lipid droplet proteins including binding via a hydrophobic face

**DOI:** 10.64898/2026.11.20.766188

**Authors:** Siqi Sun, Martin Bonin, Magdiel S. S. Lim, Philipp Niemeyer, Elena Lesch, Payton Whitehead, Kent D. Chapman, Sebastian Thallmair, Till Ischebeck

## Abstract

Plant lipid droplets (LDs) are specialized cellular subcompartments for lipid storage and key hubs for lipid metabolism. They have a unique structure: a hydrophobic core composed of neutral lipids enclosed by a phospholipid monolayer and associated proteins. Progress has been made in the identification and characterization of LD-associated plant proteins in recent years, raising the question of how these proteins bind to LDs. To investigate common structural features of LD association, we set up an *in silico* system using molecular dynamics (MD) simulations to elucidate the nature of this interaction and to identify predicted binding regions of selected proteins. We then tested the importance of these regions for LD association by transient expression of truncated or mutated variants in tobacco pollen tubes. With this combined approach, we investigated the LD targeting region of thirteen LD-associated proteins. In several proteins, a distinct structural feature, a hydrophobic face, binds to LDs by interacting with the acyl chains of the phospholipid monolayer through large hydrophobic residues thereby pushing the headgroups to the sides. We also analyzed proteins previously assayed for LD association and demonstrate a correlation between predicted binding parameters and *in vivo* binding.

**One-sentence summary:** This study investigates the molecular binding of 13 proteins to LDs through molecular dynamics simulations and *in vivo* localization studies in tobacco pollen tubes.

## Introduction

Cellular subcompartmentalization is one of the hallmarks of eukaryotic life (Brown and Doolittle, 1997). As each subcompartment requires a defined set of proteins, highly specific protein targeting mechanisms have evolved, allowing for the import of these proteins into the aqueous lumen of particular cellular subcompartments (Blobel, 1980; Rapoport et al., 2017). Other mechanisms facilitate the integration of proteins into specific membranes, or the associations with specific membrane surfaces. While the correct targeting in many cases requires helper proteins, certain proteins might interact with specific membranes directly based on defined molecular interactions (Mackenzie, 2005; Park and Rapoport, 2012; Walter and Johnson, 1994; Schleiff and Becker, 2011).

For most subcellular structures, certain amino acid stretches or motifs are known that are shared by many proteins and are essential and often sufficient for correct subcellular targeting, allowing for computational predictions based on primary sequences alone (Mackenzie, 2005). In contrast, no such sequences have been described for cytosolic plant lipid droplets (LDs). Generally, it is not well understood how LD-associated proteins recognize and bind to the LD surface.

LDs are ER-derived structures that consist of a hydrophobic core, in most cases predominantly triacylglycerol, that is surrounded by a phospholipid monolayer. The increasing number of plant LD proteins identified in recent years (Omata et al., 2024; Kretzschmar et al., 2020; Pyc et al., 2017; Kretzschmar et al., 2018; Fernández-Santos et al., 2020; Jolivet et al., 2009; Jolivet et al., 2004; Liu et al., 2015; Zaaboul et al., 2018; Zhi et al., 2017; Thakur and Bhatla, 2015; Scholz et al., 2024; Doner et al., 2021; Brocard et al., 2017; Scholz et al., 2026), paves the way to identify common targeting and binding mechanisms. The unique structure of LDs makes classical bilayer spanning protein architectures impossible. Therefore, LD proteins needed to evolve other mechanisms for specific targeting to the monolayer membrane of LDs. In the mammalian and yeast literature, three general modes of LD binding have been proposed, namely via a protein-membrane interaction, a lipid anchor, or via protein-protein interaction (Walther and Farese, 2012).

Proteins capable of binding to LDs via a protein-membrane interaction are distinguished as class I- and class II-type LD proteins. Class I encompasses proteins initially inserted into the ER membrane before being transferred to the LD surface (Kory et al., 2016) or to proteins targeting the LD monolayer via hydrophobic domains (Bersuker and Olzmann, 2017). One well-known example of a plant class I protein is oleosin, which contains a conserved central hydrophobic domain of two α-helices linked by a “proline knot” loop (Huang, 1992). Oleosin is inserted into the ER membrane co-translationally, with its N- and C-termini facing the cytosol and the hydrophobic region intercalated with the ER. After being transferred to the LD, the proline knot has been hypothesized to be inserted into the hydrophobic core of neutral lipids (Abell et al., 1997; Jolivet et al., 2017).

Class II LD proteins are defined as proteins synthesized in the cytoplasm that directly target the surface of LDs (Kory et al., 2016), for example via amphipathic helices. In plants, the mechanism for amphipathic helix targeting to the LD surface has not been well-characterized, but it has been reported in other organisms that amphipathic helices may detect (and stabilize) the neutral lipids exposed through holes in the phospholipid monolayer (termed packing defects) on the LD surface (Prévost et al., 2018; Olzmann and Carvalho, 2019). Compared with a relaxed phospholipid bilayer, the phospholipid monolayer of LDs is more prone to packing defects, which can be recognized by the side-chains of large hydrophobic amino acids in the amphipathic helix, making the LD surface a more optimal site for amphipathic helices to bind (Prévost et al., 2018; Bacle et al., 2017; Corey et al., 1993; Chorlay and Thiam, 2020). In many plant LD proteins, such as PLANT UBX-DOMAIN CONTAINING PROTEIN (PUX10), LD-ASSOICATED PROTEIN INTERACTING PROTEIN (LDIP) and SEED LIPID DROPLET PROTEIN (SLDP1), a predicted amphipathic helix next to a hydrophobic stretch comprising a total of ∼50 amino acids has been shown to be necessary and sufficient for LD targeting (Kretzschmar et al., 2018; Krawczyk et al., 2022; Pyc et al., 2017). However, it is not clear how these regions interact with the LD surface and similar motifs were not found in all identified plant LD proteins. To get further insights into the LD targeting mechanisms, we combined molecular dynamics (MD) simulations with a cell biological approach on a number of Arabidopsis LD proteins with unknown targeting mechanism and surveyed for physiochemical and structural features that may be critical for LD targeting. Here, we show that a single amphipathic/hydrophobic helix-like region is critical for LD targeting of several Arabidopsis LD proteins. Apart from that, in several known/predicted LD targeting proteins that act as enzymes with highly hydrophobic substrates, such as CALEOSIN 1 (CLO1), CYCLOARTENOL SYNTHASE 1 (CAS1), LD LIPASE (LIDL1) and LD-ASSOCIATED HYDROLASE (LDAH1), a hydrophobic face likely serves as the LD targeting site. This hydrophobic face is composed of two or more distinct amino acid stretches. We furthermore found proteins that contained several regions, each individually sufficient for targeting.

Finally, we tested if the MD simulations can be used to predict whether or not a protein can bind to the LD surface *in vivo*. Indeed, the results indicate that actual LD proteins form more close contacts with the phospholipids of the monolayer and bind in a more reproducible manner than proteins that were found not to localize to LDs.

## Materials and methods

### Arabidopsis seed LD isolation

Lipid droplet isolation protocol was adapted from Cai et al., 2015. For lipid droplet isolation from mature Arabidopsis seeds, approximately 2 mg of Col-0 seeds were imbibed in water for 15 min to soften seed coats. Seeds were then transferred to a sterilized mortar and ground with 5 ml of ice cold 600 mM sucrose buffer (600 mM sucrose, 1.25 mM EDTA, 10 mM KCl, 100mM Tris HCl, pH 7.5). Seed homogenates were then transferred to 15 ml high speed centrifuge tubes, and 2.5 ml of ice cold 400 mM sucrose buffer (400 mM sucrose, 1.25 mM EDTA, 10 mM KCl, 100mM Tris HCl, pH 7.5) was carefully layered on top. Samples were then centrifuged at 4°C at 10,500 g for 1 h in a swinging bucket centrifuge to float lipid droplets. The cloudy layer on top of the solution containing the isolated lipid droplets was then transferred to clean centrifuge tubes where a 200 mM sucrose buffer was carefully layered on top. The tubes were then centrifuged at 4°C at 10,500 g for 1 h in a swinging bucket centrifuge. The resulting isolated lipid droplets were collected, resuspended in 1 ml of the 200 mM sucrose buffer, and used for lipid extraction and subsequent lipid analysis.

### Lipid extraction lipidomic profiling

Lipid extraction and profiling protocols are taken from Bartels and Dörmann, 2021 and Romsdahl et al., 2022. For lipid extraction from isolated lipid droplets, 3.75 ml of chloroform/isopropanol (1:2 v/v, 0.01% BHT v/v) was added along with the 1 ml of isolated lipid droplets to glass test tubes and vortexed for ∼10 s. 10 µl of the Internal standard mix UltimateSPLASH ONE (Avanti Polar Lipids 330820L) was added to each sample along with 1.25 ml of chloroform and vortexed for ∼5 s. 1.25 ml of 0.9% KCL (aq) was added and vortexed for ∼5 s. Samples were then centrifuged at 1000 g for 5 min at room temperature before the bottom organic phase was transferred to fresh tubes and dried under a N_2_ stream. Samples were then resuspended in 1 ml dichloromethane:methanol (1:1 v/v) before being diluted in either 490 µl of acetonitrile:isopropanol:methanol:water (3:3:3:1 v/v/v/v) with 10mM ammonium acetate for non-polar lipid analysis or diluted in 490 µL of 100% ethanol containing 2 mM ammonium acetate for polar lipid analysis. Samples were run on instrumentation and with methods described in Romsdahl et al., 2022.

### MD simulations

The MD simulations were conducted with GROMACS 2023 (Bekker et al., 1993; Abraham et al., 2015; Berendsen et al., 1995; Hess et al., 2008; Lindahl et al., 2001; Páll et al., 2015; Pronk et al., 2013; van der Spoel et al., 2005) using the coarse-grained Martini 3 force field (Souza et al., 2021). An LD system consisting of two phospholipid (PL) layers with neutral TAGs in between with water and sodium ions on both sides was generated with Packmol (Martínez and Martínez, 2003; Martínez et al., 2009). Based on experimental data of the *A. thaliana* LD composition, the PL layers consist of 176 DFPC (simulating phosphatidylcholine [PC] with 18:3 or 16:3 acyl chains), 184 PIPC (PC 16:0/18:2), 184 POPC (PC 16:0/18:1), 64 DFPE (phosphatidylethanolamine [PE] 18:3/18:3), 72 PIPE (PE 16:0/18:2), 72 POPE (PE 16:0/18:1), 136 PIPI (phosphatidylinositol [PI] 16:0/18:2),, 136 POPI (PI 16:0/18:1) with parameters taken from Borges-Araújo et al. (Borges-Araújo et al., 2022) for PI lipids and the remaining parameters taken from the initial Martini 3 lipidome. A parameter file for 1024 TAGs, resembling glyceryl trioleate (TOG) with three C18:1 oleoyl tails, was generated manually based on DOPC according to the Martini 3 rules. The system was energy minimized using steepest descent, before running a 10 ns equilibration using the leap-frog integrator in 20 fs steps with a V-rescale thermostat at 280 K and a Parrinello-Rahman barostat at 1 bar.

Protein structures were downloaded from the AlphaFold database (Varadi et al., 2022). When displayed in the figures, the proteins are colored according to the to the Eisenberg hydrophobicity scale (Eisenberg et al., 1984). Proteins were converted to a coarse-grained structure with martinize2 if applicable using an elastic network (Souza et al., 2025; Kroon et al., 2025). The elastic bond force constant was set to 700 kJ/mol/nm² with lower and upper cutoffs of 0.5 nm and 0.9 nm, respectively. Both elastic bond decay factor and power were set to 0, and side chain corrections were applied. After conducting a short 100 step energy minimization in vacuum, the protein was randomly placed into the aqueous phase of the previously generated LD system ensuring a minimum distance of at least 2 nm between the protein and the LD system. This step was repeated ten times to generate ten independent starting structures. Counter ions for the protein were added, and the salt concentration was adjusted to 150 mM with sodium chloride. The system was energy minimized using steepest descent, before running a 1 ns equilibration using the leap-frog integrator in 20 fs steps with a V-rescale thermostat at 300 K and a C-rescale barostat at 1 bar. Finally, a 500 ns or 2000 ns production simulation was conducted using the leap-frog integrator in 20 fs steps with a V-rescale thermostat at 300 K and a Parrinello-Rahman barostat at 1 bar. A Verlet cutoff scheme was used for van der Waals interactions, and the electrostatic interactions were calculated with a reaction-field algorithm with a cut-off of 1.1 nm.

The resulting trajectories were analyzed with the GROMACS-implemented tools rms, mindist and distance. To visualize the trajectory, the protein was centered in the xy-plane while keeping the LD system stationary. These trajectories were analyzed with VMD (Humphrey et al., 1996) from which exemplary movies were generated for some proteins. Visualizations of the minimal distance between each residue and the LD system were prepared in PyMOL (Schrödinger, 2023) using the loadBfacts.py script from Pietro Gatti-Lafranconi (Pietro Gatti-Lafranconi, 2014). Additionally, further analyses of the simulations were carried out with custom scripts, which together with more detailed information about the simulations can be found in the accompanying Zenodo repository doi.org/10.5281/zenodo.15235178.

### Plasmid construction

Genes of interest were cloned into pLatmVC-GW, pLatmVN-GW, pLatmCC-GW via fast Gateway® (Thermo Fisher Scientific) as described in (Müller et al., 2017) or via Gibson Assembly® (New England Biolabs) following the manufacturer’s protocol. To mutagenize multiple amino acids of a construct, we ordered matching primers containing the desired mutations which can anneal with each other to amplify a short fragment with the desired mutations. Then, we used Gibson Assembly® to replace the corresponding sequence with the mutagenized fragment in the previously generated non-mutagenized constructs. All primers used can be found in Data S3

### Plant Growth Condition

*Nicotiana tabacum* L. cv. Samsun-NN (tobacco) plants were either grown as describe in (Kretzschmar et al., 2018), or were grown in a greenhouse under long day conditions with lights from lamps in addition to sunlight to adjust the light intensity to 230-250 μmol m^-2^ s^-1^ at leave height. The temperature was 22℃ and the relative humidity was around 65%.

### Particle bombardment and confocal laser scanning microscopy

*Nicotiana tabacum* L. cv. Samsun-NN pollen grains were transiently transformed using a gene gun, incubated on microscope slides in a humid environment, fixed and stained as described in (Müller et al., 2017).

Images were acquired by Zeiss LSM 780 or Zeiss LSM 980 confocal laser scanning microscope (Carl Zeiss). BODIPY493/503 and mVenus were excited at 488 nm. BODIPY was detected at 499-544 nm or 490-535 nm, and mVenus was detected at 499-517 nm when LDs are stained with Nile Red and detected at 490-535 nm when LDs are stained with MDH. Nile Red and mCherry were excited at 561 nm. Nile Red was detected at 580-620 nm and mCherry was detected at 597-641 nm. MDH is excited at 400 nm and detected at 419-466 nm. CFP was excited at 405 nm and detected at 455-481 nm. The major beam splitter (MBS) 405 was used for CFP and MDH, and MBS 488/561 was used for the other fluorophore/dye.

### Creation of sequence logos

The EnsemblPlants Orthologue search (EnsemblPlants Archive https://may2024-plants.ensembl.org/index.html) was queried with Arabidopsis gene IDs to retrieve alignments of orthologous protein sequences. Orthologs that strongly diverged from or entirely lacked the region(s) of interest were excluded. The refined sequence sets were then re-aligned using the MAFFT web tool (https://mafft.cbrc.jp/alignment/software/) with default parameters, and the resulting datasets were manually curated to remove remaining outliers. Finally, the alignment region spanning 10 amino acids upstream to 10 amino acids downstream of the region of interest was extracted and visualized as a sequence logo using the WebLogo online application (Crooks et al., 2004). Underlying alignments are available in Supplemental Data S4 and S5.

### Accession Numbers

CAS1, At2G07050.1; CINV1, At1G35580.1; CLO1, At4G26740.1; HSD1, At5G50600.1; LDAH1, At1G10740.1; LDDH1, At1G75180.1; LIDL1, At1G18460.1; LIME1, At4G33110.1; MYOB14, At4G13160.1; OBL1, At3G14360.1; OLE1, At4G25140.1; PUX10, At4G10790.1; SLDP1, At1G65090.3; UFAO1, At1G30130.1. Lysozyme from *Enterobacteria phage* T4, Uniprot code P00720.

## Results

### Set-up of a system for *in silico* testing of protein-LD interactions

More than twenty protein families have been identified in recent years that are associated with LDs (Guzha et al., 2023; Omata et al., 2024; Scholz et al., 2024; Dabisch et al., 2026; Scholz et al., 2026). While for some of these proteins regions were identified that play a role in targeting (Krawczyk et al., 2022; Kretzschmar et al., 2018; Pyc et al., 2017; Doner et al., 2025; Pyc et al., 2021), their exact binding mechanisms are not always clear. Furthermore, the LD binding mechanism of several additional protein families is not investigated at all.

So far, it is not possible to predict if and how proteins bind to the surface of LDs. Not all LD proteins bear a proline knot or an amphipathic helix that have been implemented in LD binding. Therefore, there might be other types of structural features involved. In order to find and investigate general mechanisms, we simulated the binding of several known LD-associated proteins (Table 1) in coarse-grained MD simulations and then tested in a cell biological approach, if the protein regions identified in the simulations were indeed important for LD binding.

**Table 1.** Overview of selected proteins.

| Protein | (Putative) function | LD binding mode |
|---|---|---|
| CALEOSIN 1.1 (CLO1.1) | Peroxygenase, stress response; | Hydrophobic face |
| CYCLOARTHENOL SYNTHASE1.1 (CAS1) | Sterol metabolism | Hydrophobic face |
| LD ASSOCIATED DEHYDROGENASE 1.1 (LDDH1.1) | Putative Enzyme | Hydrophobic helix |
| LD ASSOCIATED HYDROLASE 1.1 (LDAH1.1) | Putative TAG lipase | Amphipathic/hydrophobic helix; Hydrophobic surface |
| LD ASSOCIATED METHYLTRANSFERASE1.1 (LIME1.1) | Enzyme with putative function in furan fatty acid biosynthesis | Multiple protein segments |
| LD LIPASE1.1 (LIDL1.1) | Putative enzyme | Multiple protein segments; Hydrophobic surface |
| MYOSIN BINDING PROTEIN 1.1 (MYOB1.1) | LD-myosin association | Amphipathic/hydrophobic helix |
| OIL BODY LIPASE 1.1 (OBL 1.1) | TAG lipase | Hydrophobic surface |
| OLEOSIN 1.1 (OLE1.1) | Structural protein | Proline knot |
| PLANT UBX DOMAIN-CONTAINING PROTEIN 10.1 (PUX10.1) | LD protein degradation | Amphipathic/hydrophobic helix |
| SEED LD PROTEIN 1.3 (SLDP1.3) | Plasma membrane contact site formation | Amphipathic/hydrophobic helix |
| STEROLEOSIN 1.1 (HSD1.1) | Steroid metabolism | Amphipathic/hydrophobic helix |
| UNSATURATED FATTY ACID OXIDATION 1.1 (UFAO1.1) | Enzyme with putative function in furan fatty acid biosynthesis | Amphipathic/hydrophobic helix |

To enable the simulation of complex systems for longer time spans, we used coarse-grained MD simulation with the Martin 3 force field in which neighboring atoms are grouped and represented by beads (Souza et al., 2021). The Martini 3 force field was successfully applied to characterize membrane binding and identify lipid binding sites in peripheral membrane proteins (Thallmair et al., 2022; Lautala et al., 2026; Cheung et al., 2025; Borges-Araújo et al., 2022). Previous Martini versions as well as other coarse-grained force fields were used to study the role of seipin in LD biogenesis as well as LD-targeting proteins (Caillon et al., 2020; Kim et al., 2022). To simulate an LD, we generated a system with a 4 nm-thick TAG layer inserted between two PL monolayers (Figure 1 A). Because all candidate proteins were chosen from Arabidopsis, the PL bilayer was constructed based on the PL composition of Arabidopsis LD membranes (Table 2) that we determined by isolating LDs, extracting the lipids and analyzing them by LC-MS/MS (Table S1).

**Figure 1.**
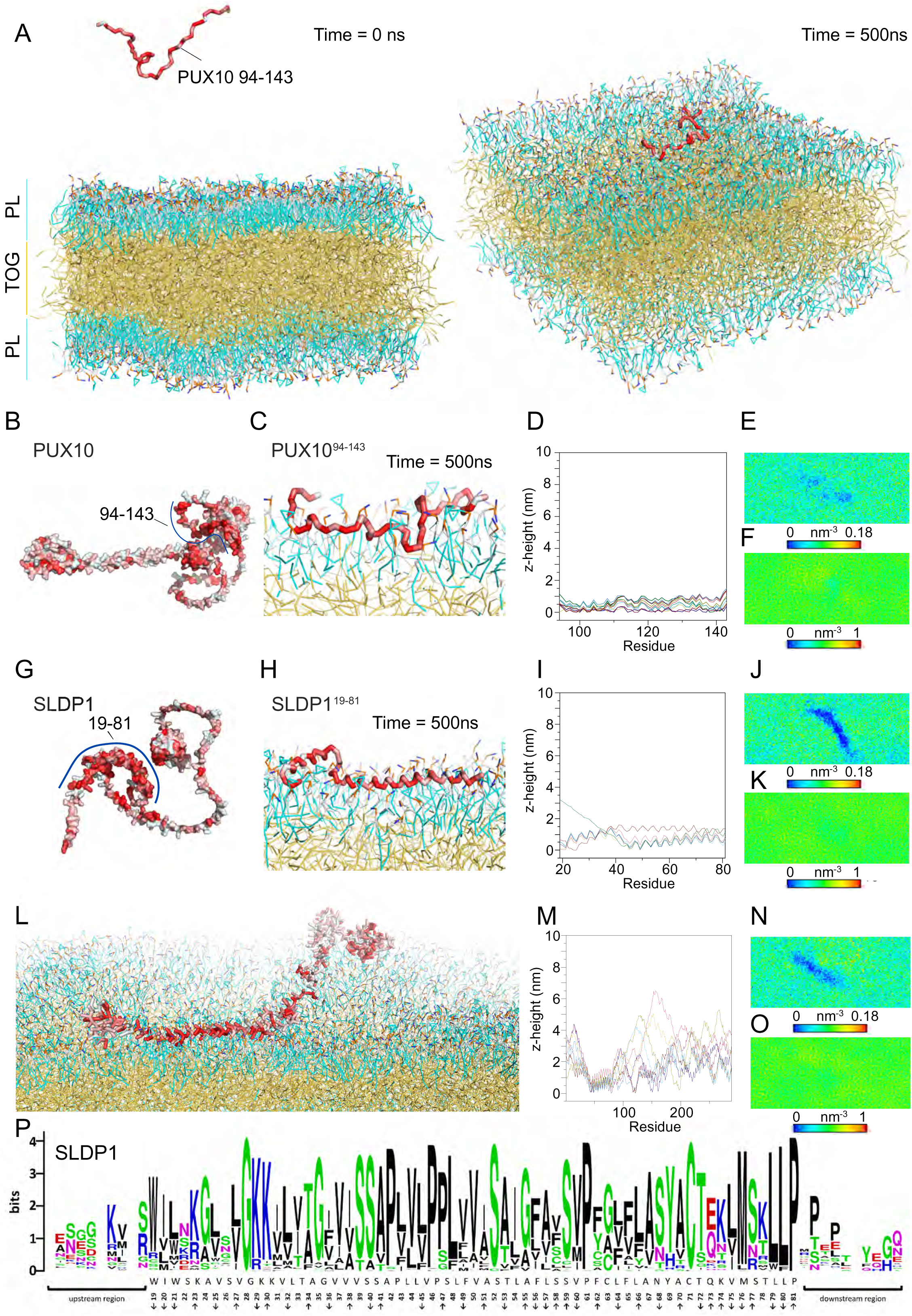
The LD targeting region of PUX10 and SLDP1 interact with the LD system in coarse-grained MD simulations. **A** Snapshot of the coarse-grained MD simulation system. The system contains the protein (here AtPUX10^94-143^), the LD system composed of PC, PE, and PI, TAG, ions, and water molecules. The protein backbone is displayed as a stick model. The ion and water molecules are not shown. Left panel, the initial state of the simulation. AtPUX10^94-143^ is placed >2 nm away from the PL monolayer in a random orientation. Right panel, last frame of the simulation in which AtPUX10^94-143^ is bound to the LD system. **B-J** AtPUX10^94-143^ and SLDP1.3^19-81^ bind to the LD system within the simulation timeframe in 10 and 7 out of 10 independent simulations, respectively. **B**, **G** AlphaFold2 models of full length AtPUX10 and SLDP1.3. The protein is shown in surface representation colored according to the Eisenberg hydrophobicity scale. Red, hydrophobic; white, hydrophilic. The reported LD targeting region is highlighted with a blue line. **C**, **H**, **L** Last frame of a representative simulation, in which the proteins bound to the LD system (time = 500 ns). **D**, **I**, **M** Per residue z-height distance of the successfully bound 10 (AtPUX10), 7 (SLDP1.3^19-81^) and 8 (SLDP1.3) independent simulations. Results from independent simulations are displayed in different colors. Averaged phosphate (**E**, **J**, **N**) and acyl chain (**F**, **K**, **O**) density of the binding leaflet in the last 100 ns of one representative simulation are displayed. **P** Sequence logo of SLDP^19-81^ and adjacent residues in an alignment of 69 homolog sequences. The orientation of the individual residues according to the simulations z-height (**I**) is displayed with arrows.

**Table 2.** LD membrane composition from isolated LDs of Arabidopsis, and used in simulations.

| **Lipid Class** | **% of lipids** | **% of lipids in simulation** |
|---|---|---|
| Phosphatidylcholine | 51.7 | 53 |
| Phosphatidylethanolamine | 19.7 | 20 |
| Phosphatidylinositol | 26.0 | 27 |
| Phosphatidylglycerol | 1.6 | - |
| Phosphatidylserine | 0.9 | - |

All protein structures were downloaded from the AlphaFold2 database (Varadi et al., 2022). Based on the individual protein structures and the prediction confidence given by the local distance difference test score (pLDDT) we used the full-length protein, a truncated version or only a specific region. For full-length proteins, we removed short apparently unstructured N- and C-terminal regions with pLDDT scores below 70 to avoid any interference from these regions during simulation. Tertiary structures were stabilized using an elastic network (Periole et al., 2009), whereas unstructured proteins or truncated versions were simulated without an elastic network. The simulation time was mainly adjusted according to the protein size. Smaller (truncated) proteins were simulated for 500 ns whereas larger proteins were simulated for 2000 ns, trying to ensure they have enough time to reach the LD system and adjust their orientation at the monolayer. All simulation data, including exemplary videos of the simulations, a detailed description of how they were performed and the necessary scripts for performing these simulations, can be found in the accompanying Zenodo repository doi.org/10.5281/zenodo.15235178.

### The reported LD targeting regions of PUX10 and SLDP1 bind to the LD surface in MD simulation

We first tested our system with the reported LD targeting regions of PUX10 (Kretzschmar et al., 2018) and SLDP1 (Krawczyk et al., 2022; Prüsener et al., 2026). Both proteins have been described to work as scaffolding proteins that interact both with LDs and other proteins (Deruyffelaere et al., 2018; Kretzschmar et al., 2018; Krawczyk et al., 2022). SLDP1 targets to LDs via a hydrophobic region (amino acids 18-81 in splicing variant 3). The LD targeting region of Arabidopsis PUX10 has not been described. However, the targeting region of *Nicotiana tabacum* PUX10 was identified corresponding to residues 94-143 of AtPUX10 (Kretzschmar et al., 2018). The homologous region of AtPUX10 tagged with the fluorophore mCherry showed co-localization with LDs when being transiently expressed in tobacco pollen tubes (Figure S1).

The predicted AlphaFold2 models of both PUX10 and SLDP1 contain large, disordered regions (Figure 1 B, G). For simplicity, we simulated only their reported LD targeting regions. SLDP1.3^19-81^ is predicted to form two connected helices, whereas AtPUX10.1^94-143^ did not show any clear secondary structure. Thus, we used an elastic network only for SLDP1.3^19-81^. The protein models were placed into the aqueous phase of the simulation system with a random orientation and a distance of at least 2 nm from the PL surface (exemplary shown for PUX10.1^94-143^ in Figure 1 A). For each protein, ten independent simulations were performed, each one with a different random initial orientation.

To follow the protein position relative to the LD system during simulation, we plotted the minimal distance between protein and LD system and the number of contacts, i.e. the number of protein-LD bead pairs that are closer than 0.6 nm to one another, over time. The resulting graphs typically show the diffusion of the protein within the aqueous phase with fluctuating distances or no contacts in the first part of the simulation. Once the protein reaches the LD system in the orientation that allows binding, the minimal distance stabilizes at a constant value, while the number of contacts increases to a constant value, often between 400-500, indicating stable binding. Occasionally, only temporal binding with a lower number of contacts or even no binding was observed throughout the simulation (Figure S2, S3). AtPUX10.1^94-143^ and SLDP1.3^19-81^ arrived at the LD system within the simulation timeframe in ten or seven out of ten independent simulations, respectively (Figure S2 A, S3 A). After binding, the proteins resided at the monolayer and the number of contacts remained stable (Figure S2 B, S3 B, see Figure 1 C, H for final binding conformation). The results indicate that known LD-binding protein domains can firmly interact with the monolayer in our MD simulation system. To display which amino acids get closest to or even insert into the LD surface, thereby indicating a function in binding, we introduced a virtual phosphate z-plane as a baseline for distance calculation. The z-plane is calculated by averaging the z-position of phosphates from the phospholipids (PLs) from the protein binding leaflet in the last 100 ns of the simulation, and the z-position of each amino acid is given by their center distance to this z-plane, named z-height. For AtPUX10.1^94-143^, and even more for the helix of bound SLDP1, the z-height patterns of independent simulations were very similar, indicating a distinct reproducible binding orientation (Figure 1 D, I, M). Furthermore, when we simulated the full-length version of SLDP1.3, the hydrophobic helices were the only region that reliably and reproducible came in close contact with the lipid monolayer (Figure 1 L, M). While in most cases large hydrophobic residues pointed towards the lipid monolayer, hydrophilic residues pointed in the opposite direction (Figure 1 P). Interestingly, these protein regions do not reach into the core of the LD but are situated close to the surface of the monolayer (Figure 1 C, H), pushing the lipid headgroups to the sides (Figure 1 E, J). Here, the density of the acyl chains of the phospholipids were not strongly affected by the insertion, indicating that the lipid tails were not strongly pushed apart (Figure 1 F, K). Together, these results indicate that the identified protein regions interact with the acyl chains of the phospholipids and maybe their headgroups but not with the triacylglycerols in the core. A sequence comparison of this AtSLDP1 region across 69 proteins from 54 angiosperm species (Figure 1 P) displayed that the LD targeting region is much stronger conserved than the adjacent regions. Especially highly conserved in this sequence are four proline residues, several positively charged residues towards the N-terminal end of the sequence and large hydrophobic residues at the C-terminal end.

### Oleosin targets the LD surface via the reported proline knot in MD simulations

We then assayed another protein, OLEOSIN 1 (OLE1), for which the targeting mechanism of the protein family is well explored, at least in yeast (Beaudoin et al., 2000). Oleosins were the first discovered LD proteins and are proposed to insert into LDs via a hydrophobic hairpin with a motif called proline knot at the tip of the hydrophobic hairpin (Loer and Herman, 1993; Huang, 1992). Experiments in yeast indicate that the proteins first insert into the ER and then into the newly forming LDs (Beaudoin et al., 2000). The originally proposed structure is remarkably similar to the structure predicted by AlphaFold2 (Figure 2 A). To avoid OLE1’s N- and C-terminal region’s hydrophobic residues binding to the hydrophobic hairpin and thus shielding it from LD interaction, we performed MD simulations of the central hydrophobic hairpin region, only (residues 33-124, Figure 2 B). In these simulations, OLE1^33-124^ bound successfully to the LD system surface in all independent simulations with a high number of contacts (Figure S4). Different from the published hypothesis, the long hydrophobic hairpin region was not inserted vertically into the monolayer, thereby reaching the hydrophobic core, but bound to the LD surface horizontally (Figure 2 B), pushing the lipid head groups aside (Figure 2 C). Amino acids 80-120 were inserted into the PL monolayer in all simulations, while the N-terminal part of the hairpin, especially residues 33-50, was often not interacting with the LD surface (Figure 2 D). Again, the orientation of these residues was reproducible, and in most cases, large hydrophobic residues pointed towards the LD surface were conserved across 81 orthologous sequences (Figure 2 E). Interestingly, regions upstream and downstream of the targeting sequence were much less conserved.

**Figure 2.**
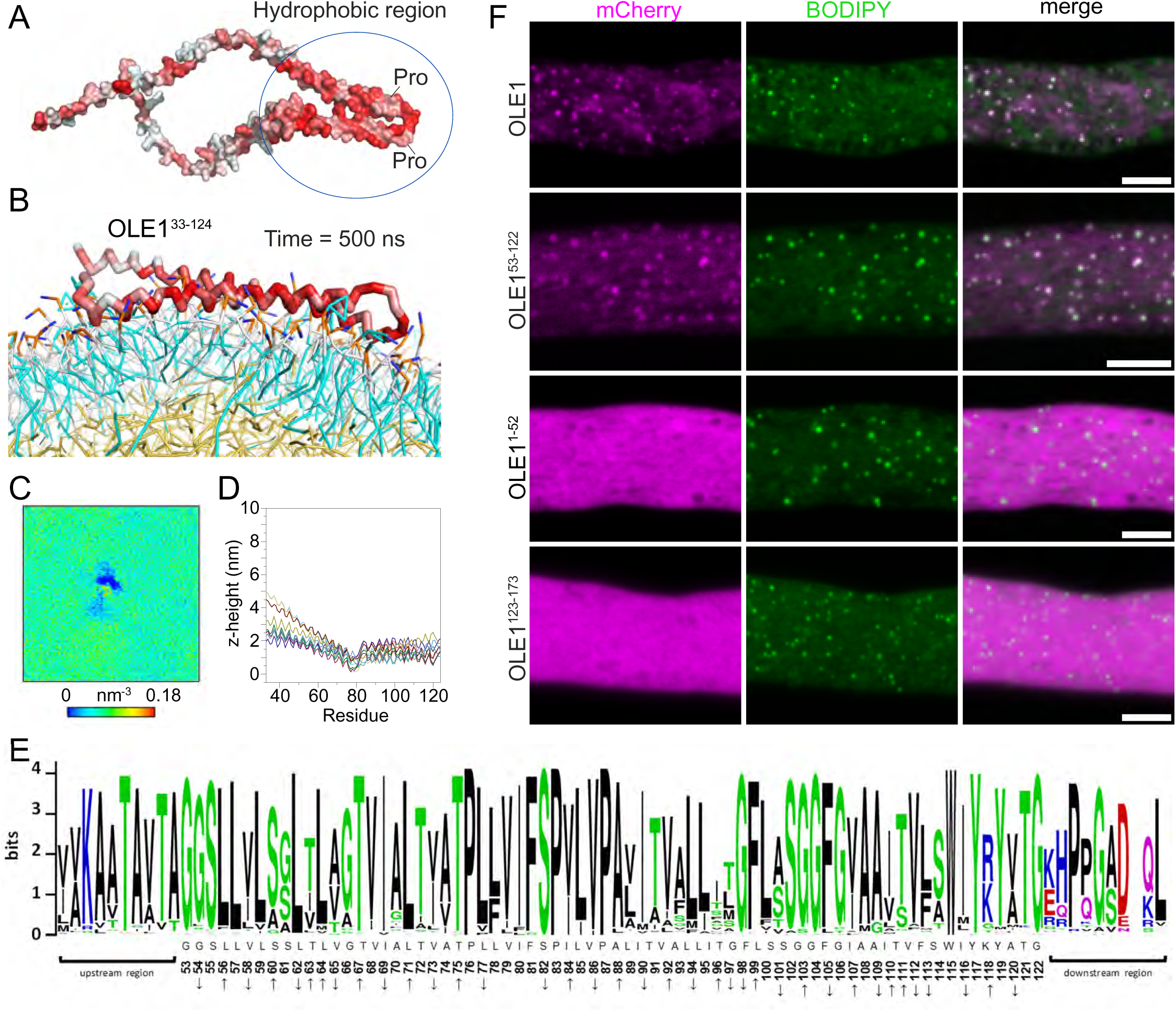
The proline knot region of OLE1 localizes to the LD surface in MD simulations and pollen tubes. **A-D** MD simulation results for OLE1. **A** AlphaFold2 model of the full-length OLE1. The annotated central hydrophobic region with proline knot can be observed. The protein is colored according to the Eisenberg hydrophobicity scale. Red, hydrophobic; white, hydrophilic. **B** Last frame of a representative simulation, in which OLE1^33-122^ bound to the LD system surface. The OLE1^33-122^ backbone is displayed as sticks. OLE1^33-122^ arrived at the LD system in 10 independent simulations during the simulation timeframe (time = 500 ns). **C** First panel, phosphate density plot of one representative MD simulation of OLE1^33-124^. **D** Per residue z-height result of 10 independent simulations. Results from independent simulations are shown in different colors. **E** Sequence logo of OLE1^53-122^ and adjacent residues in an alignment of 81 homolog sequences. The orientation of the individual residues according to the simulations (**D**) is displayed with arrows. **F** Subcellular localization of OLE1 and its truncated versions. The full length OLE1 and OLE1^53-122^ co-localized with LDs, whereas the N-terminus and C-terminus of OLE1 were cytosolic. All constructs were fused with mCherry at the C-terminus (magenta channel). Pollen tubes were incubated for 5-7 h before imaging. LDs were stained with BODIPY 493/503 (green channel). Scale bars, 5 µm.

Based on these findings, we cloned a sequence encoding a truncated version of the central hydrophobic domain with a shorter N-terminal region, OLE1^53-122^, for expression in pollen tubes. This truncated version showed clear co-localization with LDs, whereas OLE1^1-52^ and OLE1^123-173^, missing the central hydrophobic domain were cytosolic (Figure 2 F).

### Soluble proteins did not bind to the LD surface in a reproducible manner in MD simulations

We also tested our system for two soluble proteins, lysozyme from *Enterobacteria phage* T4 (Uniprot code P00720) and CYTOSOLIC INVERTASE 1 from Arabidopsis (Uniprot code Q9LQF2). Although these proteins came into contact with the membrane in some of the simulations (Figure S5 A, S6 A), they could not bind properly, indicated by the number of contacts which remained much lower than for the previously tested proteins (Figure S2-S6 B). Furthermore, the simulated interaction was most often only transient and the orientation at the end of the simulation was not reproducible (Figure S5 C, S6 C). In conclusion, these results indicate that our system is able to distinguish between proteins that can potentially bind LDs and non-LD-binding proteins allowing for potential insights into binding orientation of LD proteins to be gained.

### Several proteins bind to LDs via an amphipathic/hydrophobic α-helix

After testing our system for known LD targeting protein regions and two hydrophilic proteins, we assayed it for LD-associated proteins (Kretzschmar et al., 2020; Lin et al., 2002; Omata et al., 2024; Qu et al., 1986; Chen et al., 1999) for which neither the binding region nor binding orientation is known. Whereas different replicates of the ten simulations performed per protein might result in different binding modes, we hypothesize that the number of contacts can help to distinguish between the natural binding state and less stable binding orientations. We also hypothesize that the natural binding state dominates in the majority of simulations. Based on these presumptions, we aim to decipher if, and with which parts, the proteins bind to and how they interact with the LD surface.

According to the putative LD targeting, structural features observed in the AlphaFold2 models, MD simulation results, and results from localization assays, we classified the investigated proteins into three groups, i.e. proteins targeting to LDs via a helix, proteins targeting to LDs via a hydrophobic surface region, and proteins with multiple LD-targeting regions.

Among the investigated proteins, we found several to target LDs via an amphipathic or hydrophobic helix, including MYOSIN BINDING PROTEIN 14 (MYOB14), LD-ASSOCIATED DEHYDROGENASE 1 (LDDH1), HYDROXYSTEROID DEHYDROGENASE 1 (HSD1), and UNSATURATED FATTY ACID OXIDATION 1 (UFAO1).

The first protein we analyzed, MYOB14 is a recently identified LD protein that can interact with myosin XIK (Omata et al., 2024). When testing the full-length protein (Figure 3 A) in our system, we found that it bound to the LD via its N-terminal region in 8 of 10 simulations, pushing the head groups of the monolayer to the side (Figure 3 B, C). According to the z-height, amino acids 1-48 and in several cases also amino acids 54-85 were in closest contact with the monolayer (Figure 3 D). According to the AlphaFold2 model, the first 48 amino acids fold into an amphipathic/hydrophobic helix. As this region (MYOB14^1-50^) bound the surface in the most reliable manner, we also tested if it alone binds to the surface in MD simulations, which was indeed the case (Figure S7). We then transiently expressed mCherry-tagged MYOB14^1-50^and its complementary truncated variant mCherry-MYOB14^51-282^ in tobacco pollen tubes. MYOB14^1-50^ showed clear LD co-localization, whereas MYOB14^51-282^ was distributed in the cytosol (Figure 3 E), suggesting that MYOB14^1-50^ is necessary and sufficient for LD targeting.

**Figure 3.**
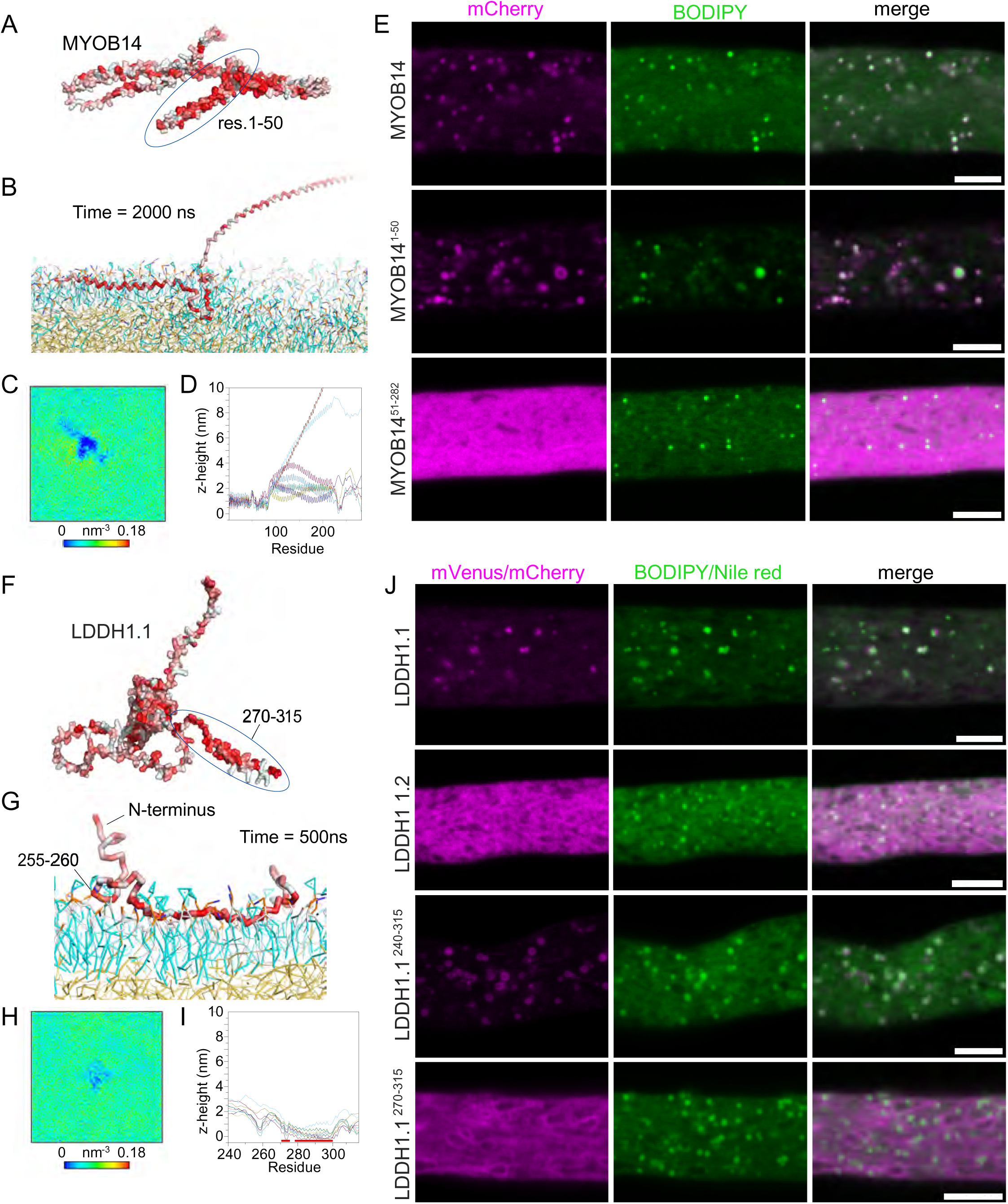
MYOB14 and LDDH1 bind to LDs in MD simulations and in tobacco pollen tubes only via a short terminal region. **A**, **F** AlphaFold2 models of MYOB14 and LDDH1.1 displayed in surface representation and colored according to the Eisenberg hydrophobicity scale. Red, hydrophobic; white, hydrophilic. Encircled in blue are amphipathic/hydrophobic helices or regions. MYOB14 and LDDH1.1 arrived at the LD system in 8 and 9 out of 10 independent simulations during the simulation timeframe (time = 2000 and 500 ns, respectively). **B**, **G** Last frames of representative simulations, in which the proteins bound to the LD system. The protein backbone is displayed as sticks. **C**, **H** Averaged phosphate density of the binding leaflet in the last 100 ns of one representative simulation. Both proteins pushed the PL head groups at the protein binding site away, showing that it was embedded into the PL monolayer in the simulations. **D, I** Per residue z-height distance of the successfully bound simulations. Results from independent simulations are shown in different colors. **E** Subcellular localization of MYOB14 and its truncated versions in tobacco pollen tubes. Pollen tubes were transiently transformed with full-length MYOB14-mCherry, MYOB14^1-50^-mCherry or MYOB14^51-282^-mCherry. LDs were stained with BODIPY 493/503. **J** LDDH1.1 and its truncated/splicing variant were fused with mVenus or mCherry (magenta channel) and the LDs were stained with Nile Red or BODIPY 493/503 (green channel), respectively. LDDH1.1 and LDDH1.2 were tagged at the C-terminus. In contrast, the C-terminal truncated versions, LDDH1.1^240-315^ and LDDH1.1^270-315^ carried the tag at the N-terminus. All pollen tubes were incubated around 5-7 h before imaging. Scale bars, 5 µm.

The next protein investigated, LDDH1, has a globular domain flanked by disordered regions and a highly hydrophobic C-terminal α-helix according to the predicted model (Figure 3 F, Figure S 8). This protein was reported to target the plastid outer envelope in tobacco BY-2 cells in a tail-anchored fashion with the amino acids 270-315 being sufficient for targeting (Teresinski et al., 2019). In *N. tabacum* pollen tubes and *N. benthamiana* leaves, the full-length version was shown to bind LDs instead (Kretzschmar et al., 2020).

When simulating full length LDDH1 with its large, disordered regions, the flexibility of the protein caused binding in many different modes, which in the simulated 2000 ns did not converge to a common distinct binding mode (see Zenodo repository doi.org/10.5281/zenodo.15235178 for simulation results). Therefore, we simulated only the C-terminus (LDDH1^240-315^) which bound to the LD surface in nine of ten independent simulations. The residues 255-260 and 270-300 were in close contact with the PL monolayer when LD binding was successful (Figure 3 G-I). The orientation of the individual amino acid residues in these regions was conserved in most of the simulations.

As the simulations and the previously published information (Teresinski et al., 2019), indicated binding via the C-terminus, we tested this hypothesis *in vivo* using several truncated variants of LDDH1 (Figure 3 J). LDDH1 also has one splicing variant, LDDH1.2, which is missing residues 240-315 of LDDH1.1, and was assessed *in vivo* as well. When transiently expressed in tobacco pollen tubes, LDDH1.2-mVenus was cytosolic. We then tested LDDH1^270-315^ that was previously shown to target the plastid envelope. Interestingly, this variant did not co-localize with LDs but with a structure reminiscent of the ER instead. As residues 255-260 seemed to play a role in binding in our simulations, we also assayed a less truncated variant, mVenus-LDDH1.1^240-315^. In comparison to LDDH1^270-315^, this construct displayed clear LD targeting.

HSD1 and UFAO1 are predicted or shown to be enzymes. HSD1 has 11β-, 17β-hydroxysteroid dehydrogenase activity (d’Andréa et al., 2007) and has been reported to be involved in brassinosteroid synthesis (Li et al., 2007; Baud et al., 2009), and UFAO1 is homologous to a bacterial enzyme that is involved in furan-containing fatty acid synthesis (Omata et al., 2024). HSD1 possesses an N-terminal region, which was hypothesized to form a hairpin structure with a proline knob at the turn (Lin et al., 2002). However, in the AlphaFold2 model, the N-terminus of HSD1 did not form such a short hydrophobic hairpin but rather an N-terminal potential amphipathic/hydrophobic helix linked to a globular domain at the C-terminus (Figure 4 A). UFAO1 displayed a similar structure (Figure 4 I). In MD simulations, the full-length HSD1 and UFAO1 successfully bound to the LD system in 5 and 8 simulations, respectively (Figure 4 B, J). The N-terminal helices of both proteins were embedded in the PL monolayer if the binding was successful (Figure 4 C, K). When expressed in tobacco pollen tubes, HSD1^1-47^ and UFAO1^1-50^ showed dual localization to LDs and the ER (Figure 4M-N, Figure S9), whereas the complementary N-terminally truncated variants (HSDx, UFAO1y) lost their ability for surface binding and were distributed throughout the cytosol. These results suggest that the N-terminal helices of HSD1 and UFAO1 are necessary for membrane association, but that other parts of the protein are important to favor LD rather than ER localization.

**Figure 4.**
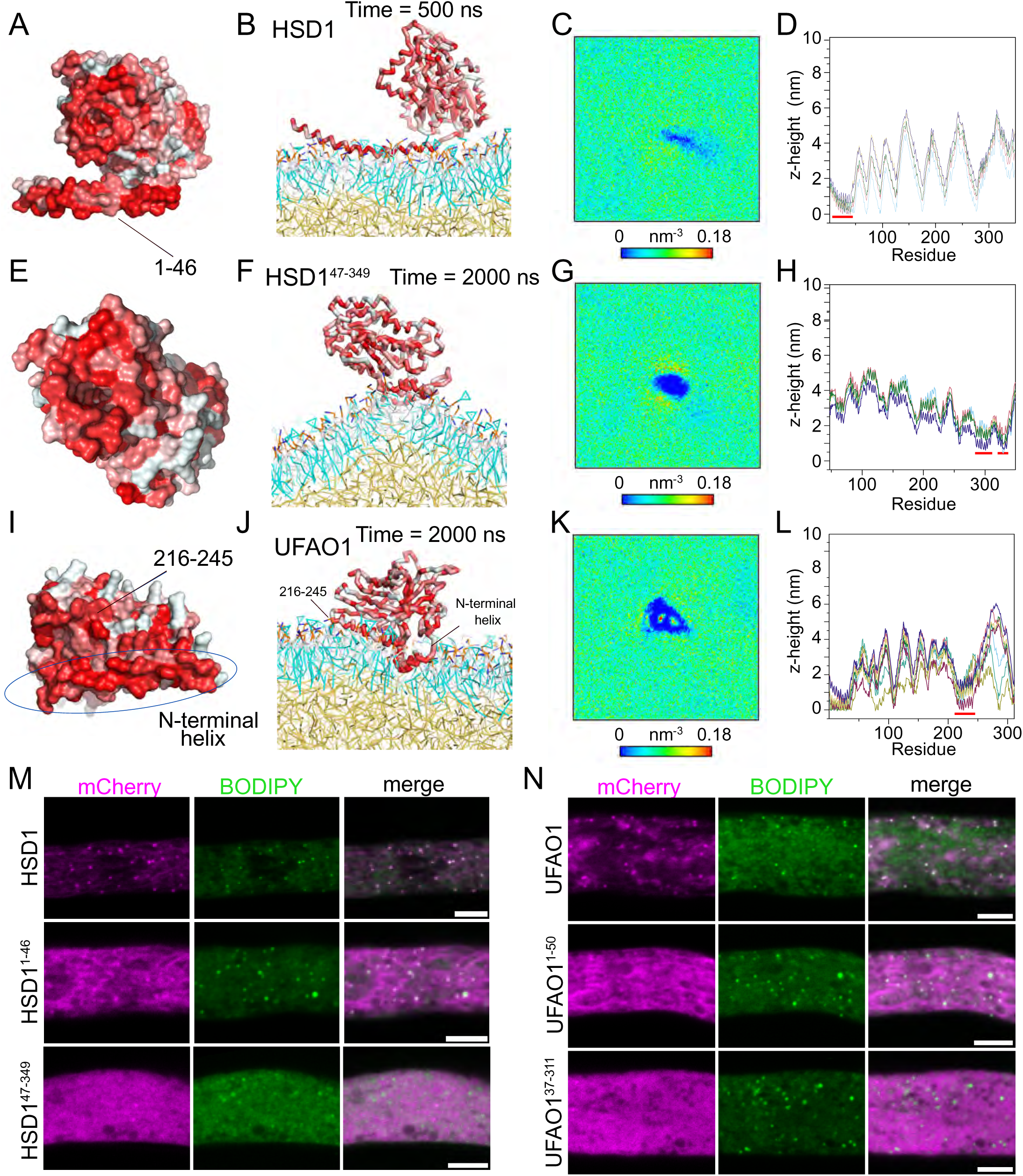
HSD1 and UFAO1 require the N-terminal amphipathic/hydrophobic helix for LD targeting. **A**, **E**, **I** AlphaFold2 models of HSD1, HSD1^47-349^ and UFAO1. Proteins are colored according to Eisenberg hydrophobicity scale. Red, hydrophobic; white, hydrophilic. **B**, **F**, **J** Last frames of representative MD simulations in which the proteins bound to the LD system. The protein backbone is displayed as sticks. If the LD binding was successful, the N-terminal helix of HSD1 and UFAO1 were interacting with the PL monolayer. UFAO1, HSD1 and HSD1^49-349^ arrived successfully at the LD system surface in 5, 5 and 8 independent simulations within the indicated simulation timeframe. **C**, **G**, **K** Phosphate density map of the binding leaflet in the last 100 ns of one representative simulation. **D**, **H**, **L** Per residue z-height distance of the successfully bound simulations. Results from independent simulations are shown in different colors. The LD system interacting region is highlighted with red lines. **M**, **N** Subcellular localization of HSD1, UFAO1, and their truncated versions in tobacco pollen tubes. All proteins were tagged with mCherry at the C-terminus. The pollen tubes were incubated around 5-7 h before imaging. LDs were stained with BODIPY 493/503. Scale bars, 5 µm.

According to the AlphaFold2 model and MD simulation results, both proteins have additional hydrophobic surface regions in their C-terminal domains (Figure 4A, E, I). For UFAO1, this hydrophobic face is structurally close to the N-terminal helix, and both these structural features participated in surface contacts with the LD system in our MD simulations (Figure 4 J-L). For HSD1, the C-terminal domain contains two hydrophobic surface regions. The first hydrophobic region consists of two helices, residues 150-170 and 194-214 and the second hydrophobic region covers residues 281-335. Neither of them interacts with the PL surface when the full-length protein was simulated. However, in MD simulations of HSD1^47-349^, the truncated protein bound to the LD system in five of ten simulations, and the binding was facilitated through the hydrophobic surface residues 281-335 (Figure 4H). We attribute this difference to the elastic network, which keeps the N-terminal helix in a fixed position towards the globular domain preventing it from adapting a conformation which allows simultaneous binding of the helix and the hydrophobic surface. Considering the full-length UFAO1 and HSD1 showed clearer LD localization *in vivo* compared with their N-terminal helices (Figure 4 M, N), it is likely that the hydrophobic surface regions in their C-terminal domains contribute to LD targeting.

### Proteins can bind to LDs via a hydrophobic face

While amphipathic helices and hairpins seem to be recurring structures important for LD targeting, they are not present in all identified LD proteins. Among our selected proteins, the AlphaFold2 models of CYCLOARTENOL SYNTHASE 1 (CAS1), CALEOSIN 1 (CLO1) and OIL BODY LIPASE 1 (OBL1) did not show any protruding helix/hairpin (Figures 5-6). Instead, the structures are spherical and have a hydrophobic region on one surface that we refer to as a hydrophobic face.

**Figure 5.**
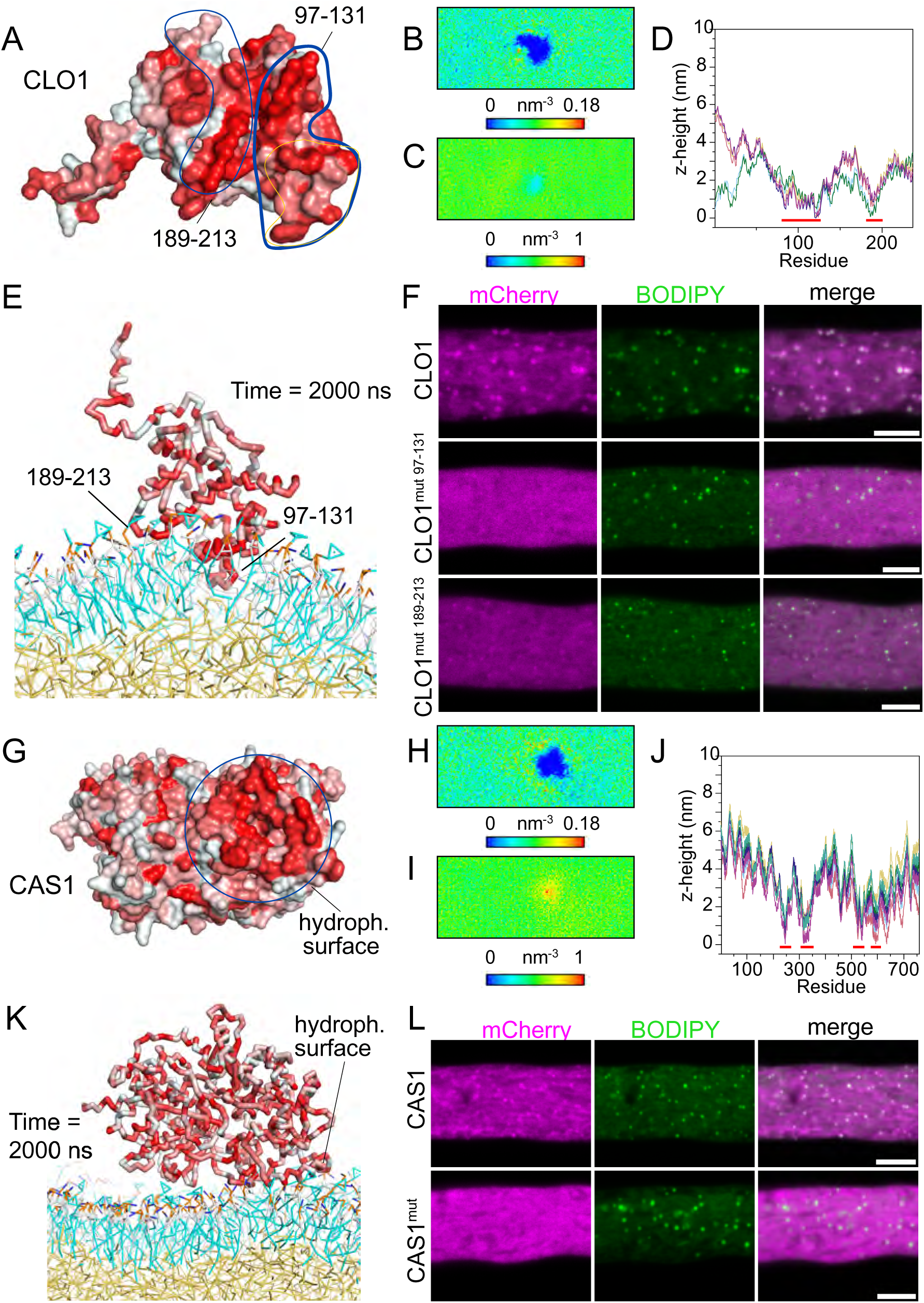
CLO1 and CAS1 interact with the LD surface through a hydrophobic region on the surface. **A**, **G** Surface representations of CLO1 and CAS1 AlphaFold2 models. The hydrophobic surface regions are highlighted by blue outlines. The proteins are colored according to the Eisenberg hydrophobicity scale. Red, hydrophobic; white, hydrophilic. **E**, **K** Last frame of a representative simulation, in which the proteins bind to the LD system surface. The protein backbones are shown as sticks. CLO1 and CAS1 arrived successfully at the LD system surface in 6 and 7 independent simulations, respectively. **B**, **H** Phosphate density plots of one representative simulation. **C**, **I** Acyl chain density plot of one representative simulation. **D**, **J** Per residue z-height results of simulations where binding was completed. The LD binding regions are highlighted with red underlines. Results from independent simulations are shown in different colors. **F**, **L** The subcellular localization of wild-type and mutagenized CLO1 and CAS1 (See Supplemental Table S1 for details). The proteins were tagged with mCherry at their C-terminus and expressed in pollen tubes. LDs were stained with BODIPY 493/503 (green panel). Scale bars, 5 µm.

**Figure 6.**
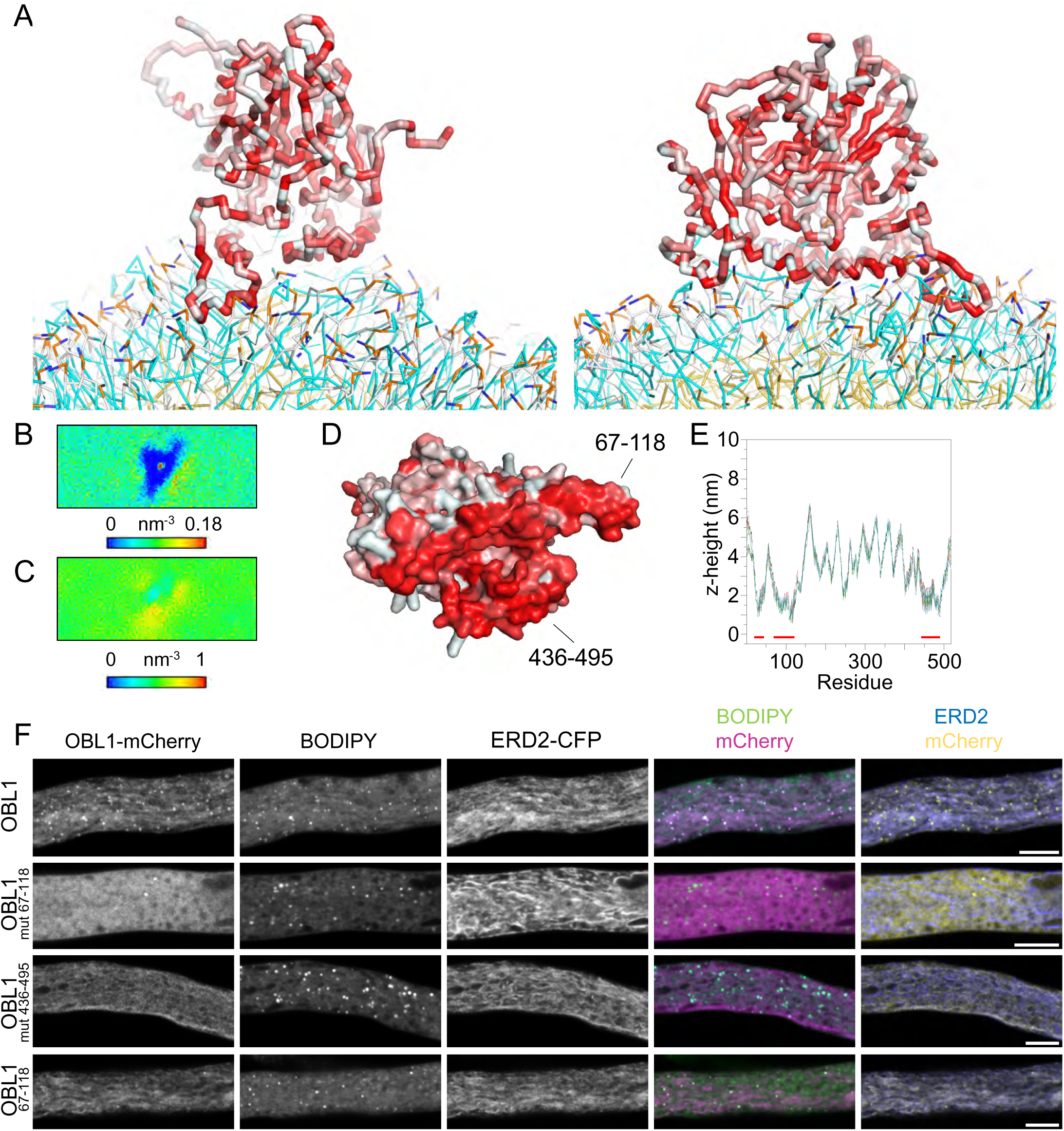
OBL1 interacts with the LD surface through a hydrophobic face. **A** Last frame of a representative simulation shown from two sides, in which the protein binds to the LD system. The protein backbone is shown as sticks. OBL1 arrived successfully at the LD system surface in 7 of 10 independent simulations. The protein is colored according to the Eisenberg hydrophobicity scale. Red, hydrophobic; white, hydrophilic. **B** Phosphate density plot of one representative simulation. **C** Acyl chain density plot of one representative simulation. **D** AlphaFold2 model displayed in surface representation. The two LD binding sections are highlighted. **E** Per residue z-height results of simulations where binding was completed. The LD binding regions are highlighted with red underlines. **F** The subcellular localization of wild-type and mutagenized (See Supplemental Table S1 for details) or truncated regions. The proteins were tagged with mCherry at their C-termini and expressed in pollen tubes and co-expressed with the ER marker ERD2-CFP. LDs were stained with BODIPY 493/503. Scale bars, 5 µm.

In previous studies, CLO1 was proposed to target LDs via a proline knot motif within a hydrophobic hairpin similar to oleosin (Chen et al., 1999). However, in the AlphaFold2 model, CLO1 was a globular protein (Figure 5A), and the predicted proline knot (residues 118-127) was not stretched out but is part of a hydrophobic face. This surface is composed of two helices, residues 97-131 and 189-213. In MD simulations, CLO1 successfully localized to the LD surface in six out of ten independent simulations (Figure 5 E). When the binding was successful and the number of contacts reached ∼300 (Figure S10A), both helices 97-131 and 189-213 interacted with the PL monolayer (Figure 5 D), pushing the phospholipid head groups and to a lower extent even the phospholipids acyl chains at the binding site outwards (Figure 5 B, C). Therefore, we hypothesized that CLO1 might target LDs via this hydrophobic face.

Compared with a proline knot or an amphipathic helix, maintenance of a hydrophobic surface proposedly relies on the intact protein structure. Thus, to avoid interrupting the tertiary structure by removing parts of the protein, we generated two full length constructs with mutagenized CLO1: (I) nine surface exposed large hydrophobic residues (F, W, I, L, Y and M) between residues 97-131, and (ii) six large hydrophobic residues between 189-213 were selected and mutagenized to valine, respectively (Table S1). These exchanges keep the area hydrophobic while removing large hydrophobic residues that were proposed to insert into packing defects of the LD surface (Prévost et al., 2018). CLO1^mut^ ^97-131^ lost its membrane targeting ability and became cytosolic, and CLO1^mut189-213^ showed only weak LD co-localization (Figure 5F). These results suggest that the hydrophobic surface is necessary for LD targeting, and that the abundance of large hydrophobic amino acid residues is critical for LD binding efficiency.

By a similar approach, we investigated the LD targeting regions of CAS1 and OBL1 (Figures 5 G-L and 6). In the simulations, likewise to CLO1, the hydrophobic faces of CAS1 and OBL1 pushed the phospholipid headgroups to the side to interact with the acyl chains of these lipids. Unlike CLO1, which pushed the lipid tails aside, CAS1 and OBL1 simulations showed an increased acyl chain density in some regions of the binding site, indicating that the acyl chains are attracted by the hydrophobic face (Figure 5 H, I, K and 6 A-C).

CAS1 targeted the LD surface via the hydrophobic surface region in seven of ten simulations (Figure 5 K) through ∼300 contacts (Figure S10B). After mutagenizing six large hydrophobic amino acid residues within this surface region to alanine (See Table S1 for details), CAS1 did not target LDs anymore when expressed in pollen tubes (Figure 5 L) but overlapped partially with an ER marker (Figure S9).

For OBL1, the number of contacts was even higher (∼600, Figure S10 C) in the seven simulations in which the protein bound via its hydrophobic face (Figure 6 A). In OBL1, the hydrophobic face comprises two amino acid stretches, 67-118 and 436-495 (Figure 6 D).Mutagenesis of 20 large hydrophobic amino acids to valine in the first region 67-118 led to a cytosolic localization, while similar mutagenesis of 12 amino acids in the second region 436-495resulted in ER-targeting (Figure 6 F). As this indicated that the first region alone might be able to target the protein to the ER, we cloned this short region attached to a C-terminal fluorophore, which indeed localized to the ER.

In conclusion, apart from the classical proline knot or amphipathic/hydrophobic helix, an LD protein may target LDs via a (to our knowledge) previously undescribed feature, a hydrophobic face. The amino acids with large hydrophobic side chains on the surface region seem to play critical roles in LD binding efficiency.

### LIME1 does not interact strongly with the monolayer in MD simulations

LIME1 residues 79-350 are annotated as an S-adenosyl-L-methionine-dependent methyltransferase superfamily by InterPro (Blum et al., 2025) and have been proposed to be involved in the in furan-containing fatty acid synthesis in Arabidopsis (Omata et al., 2024). Different from the other selected proteins, LIME1 indeed got close to the LD surface in all the 10 independent simulations (Figure 7 A). However, the phosphate density of the protein-binding PL monolayer (Figure 7 B) was influenced opposite to what was observed for all other proteins (Figures 1-6). Whereas the other proteins pushed the PL head groups aside while inserting into the PL layer, LIME1 seems to attract these head groups creating an area of higher phosphate density below the protein. Furthermore, no common orientation pattern was observed (Figure 7 C) and the number of contacts was much lower than in any other LD-binding protein (Figure S11). These results suggest that LIME1 was not inserted into the PL monolayer during the 2000 ns simulations but instead interacted with the PL head groups and pulled them towards the protein. Without a good hint from the MD simulations, we analyzed the subcellular localization of a series of LIME1 variants (Figure 7 F). Neither the N-terminal region alone (mVenus-LIME1^1-48^), nor the variants without it, namely mVenus-LIME1^291-355^, LIME1^49-290^-mVenus and LIME1^49-355^-mCherry, targeted to LDs. LIME1^1-290^-mCherrry targeted to LDs in 6 out of 10 randomly selected transformed pollen tubes. Notably, the recombinant protein, LIME1^1-48^-mVenus-LIME1^290-355^, with the mVenus being situated in between the N- and C-terminal regions, showed clear co-localization with LDs. These results demonstrate that the N-terminus of LIME1 is necessary for LD targeting, and that it might require the C-terminus to better define their subcellular localization although it remains unclear whether this LD targeting is a direct interaction with the LDs surface or an indirect interaction via protein-protein interactions.

**Figure 7.**
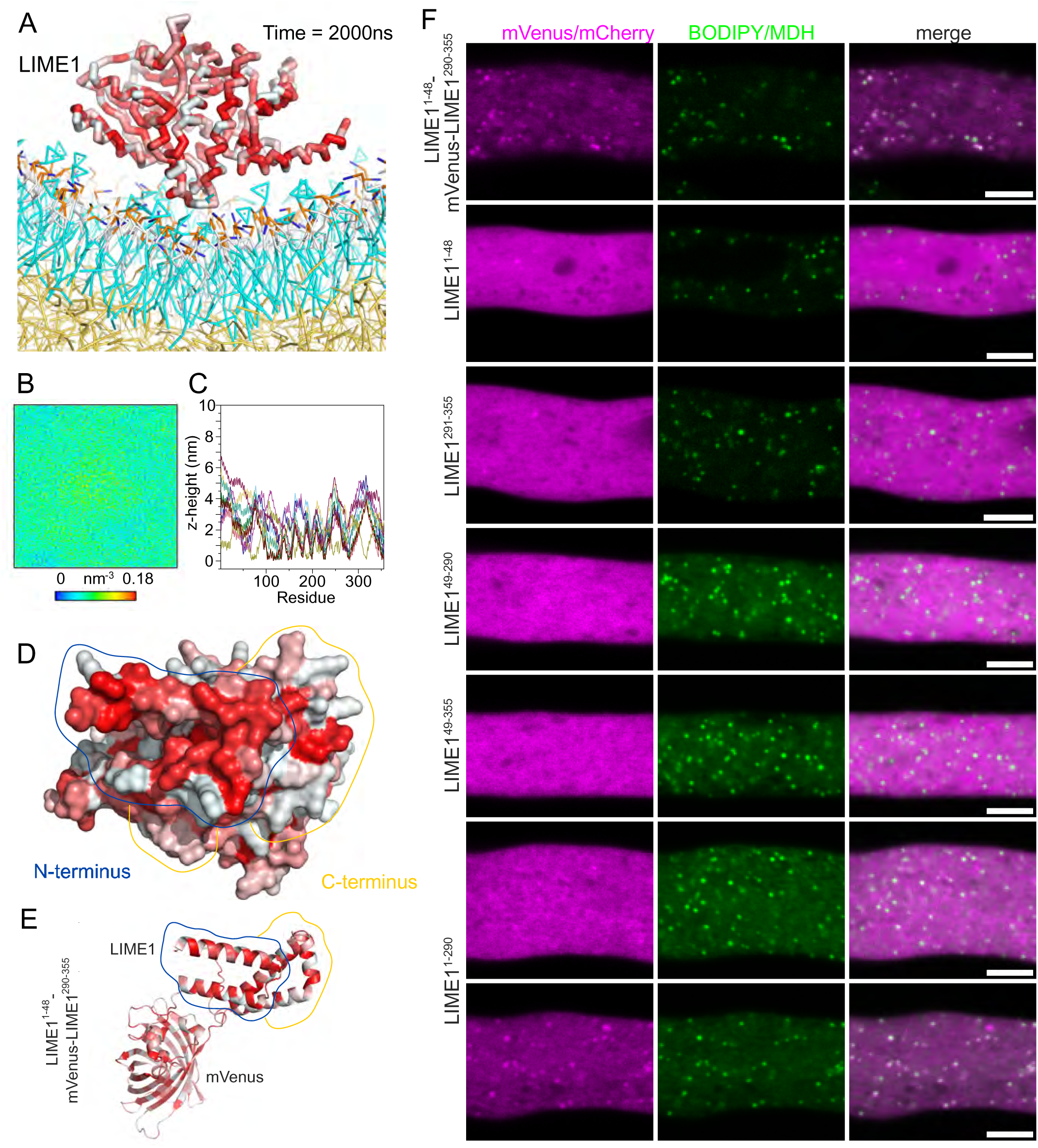
LIME1 requires multiple protein segments for LD targeting but does not have a conserved binding orientation in MD simulations. **A** Last frame of one representative MD simulation. The LIME1 backbone is shown as sticks. The protein is colored according to the Eisenberg hydrophobicity scale. Red, hydrophobic; white, hydrophilic. **B** Phosphate density plot of one representative MD simulation. **C** Per residue z-height result of 10 independent simulations. Results from independent simulations are shown in different colors. **D** AlphaFold2 model of LIME1 showed that LIME1^1-48^ (marked with a blue line) formed a small hydrophobic region on the surface that is structurally supported by the C-terminus, LIME1^290-355^ (marked with a yellow line). **E** AlphaFold3 prediction of LIME^1-48^-mVenus-LIME1^290-355^. **F** Subcellular localization of LIME1 constructs in tobacco pollen tube. LIME1^1-290^ co-localized with LDs in 6 out of 10 pollen tubes, whereas LIME1^1-48^-mVenus-LIME1^290-355^ localized to LDs in 12 out of 14 pollen tubes. Fusion proteins without LIME1^1-48^ were all cytosolic. For LIME1^1-48^, LIME1^49-355^ and LIME1^290-355^, representative images of 10 pollen tubes were shown. The pollen tubes were incubated for 5-7 h and stained with BODIPY 493/503 (for proteins tagged with mCherry) or MDH (monodansylpentane, for proteins tagged with mVenus) before imaging. Scale bars, 5 µm.

### Proteins can harbor multiple LD targeting sites

In the previous sections, we described proteins targeting to the LD surface predominantly via a single structural feature or a combination of two features necessary for sufficient binding. Interestingly, LD-ASSOCIATED HYDROLASE 1 (LDAH1) and LD lipase 1 (LIDL1) have two LD targeting sites, each of them alone being sufficient for LD targeting.

The model of LDAH1.1 shares many similarities with HSD1 and UFAO1. It has a partially hydrophobic helix of roughly 50 amino acids at the N-terminus, which is connected to a globular domain (Figure 8 A). On one side of this globular domain, a large hydrophobic face can be observed. The C-terminus of LDAH1.1 (residues 444-475) is predicted to be a disordered region and was thus removed for the MD simulations resulting in simulations of LDAH1.1^1-443^, LDAH1.1^1-50^ and LDAH1.1^67-443^ (Figure 8 B, F, J). These variants arrived at the LD surface in 7, 9 and 7 independent replicates, respectively. In all cases, the proteins displaced the head groups of the phospholipids (Figure 8 C, G, K). Both the amphipathic/hydrophobic helix and the hydrophobic face interacted with the LD surface when LD binding was successful (Figure 8 D, H, L). When being transiently expressed in tobacco pollen tubes, LDAH1.1^1-50^ showed dual localization to both the ER and the LDs (Figure 8M, S9). LDAH1.1^51-473^, on the other hand, co-localized with LDs but not with the ER (Figure 8M). Similar to CLO1 and CAS1, we selected 10 large hydrophobic amino acids within the hydrophobic face and mutagenized them to valine. The mutagenized LDAH1.1^51-473^ lost its LD co-localization and became cytosolic. Moreover, the splicing variant LDAH1.2 missing the residues 406-473 co-localized with LDs, indicating that this C-terminal region is not important for LD targeting (Figure S12). The results suggest that LDAH1 has two independent LD targeting sites, a hydrophobic helix and a hydrophobic face.

**Figure 8.**
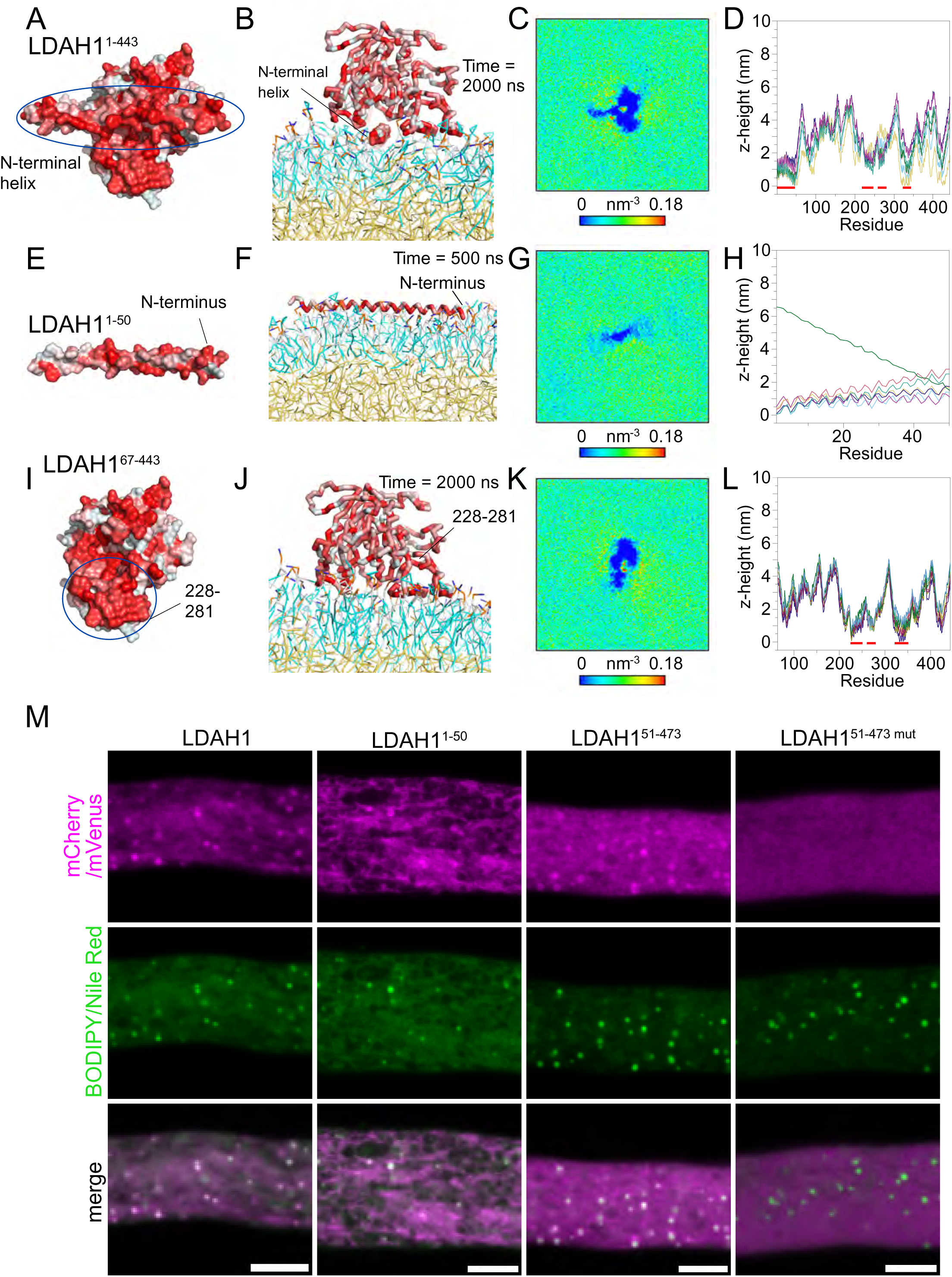
LDAH1 has two LD targeting regions that can work independently. **A-L** MD simulation results of LDAH1.1 (**A**-**D**), LDAH1.1^1-50^ (**E**-**H**) and LDAH1.1^67-443^ (**I**-**L**). **A**, **E**, **I** AlphaFold2 models of the proteins in surface representation colored according to the Eisenberg hydrophobicity scale. Red, hydrophobic; white, hydrophilic. **B**, **F**, **J** Last frame of a representative simulation, in which the protein or its truncated version bound to the LD system surface. The protein backbones are shown as sticks. LDAH1.1, LDAH1.1^1-50^ and LDAH1.1^67-443^ successfully bound to the LD system surface in 7, 9 and 7 out of 10 independent simulations, respectively, within the indicated simulation timeframe. **C**, **G**, **K** Phosphate density plots of one representative MD simulation. **D**, **H**, **L** Per residue z-height result of the successfully bound simulations. The LD system interacting regions are highlighted with red lines. Results from independent simulations are shown in different colors. **M** The full-length LDAH1.1 and LDAH1.1^1-50^ targeted to LDs and to structures looking similar to the ER. LDAH1.1^51-473^ co-localized with LDs but did not target to ER-like structures. The mutagenized LDAH1.1^51-473^ became cytosolic. Pollen tubes were incubated for 6 h before imaging. Transformation, fixation and imaging of LDAH1.1^51-473^ and its mutant were performed back-to-back. For all the constructs, the fluorescent tag was fused to the C-terminus of the protein. LDs were stained with BODIPY 493/503 when the protein was fused with mCherry and with Nile Red when the fluorophore was mVenus. Scale bars, 5 µm.

The second protein with multiple independent LD targeting regions, LIDL1, is annotated with an Alpha/Beta hydrolase fold from residues 292-670 by InterPro (Blum et al., 2025) (Figure 9A). The N-terminal region of the LIDL1 AlphaFold2 model consists of several helices, including a hydrophobic helix from residues 135-158 and an amphipathic helix from residues 205-232, spaced out by disordered regions (Figure 9B). To avoid simulating this large flexible region of the protein, we performed MD simulations of LIDL1^291-676^, which contains the putative catalytic domain. LIDL1^291-676^ bound to the LD surface in 6 independent simulations, interacting through a hydrophobic face (Figure 9 C). The protein came in contact with the monolayer, especially with residues 480-600 in a reproducible orientation. (Figure 9 D) displacing the phospholipid head groups in a ring-like pattern (Figure 9 E).

**Figure 9.**
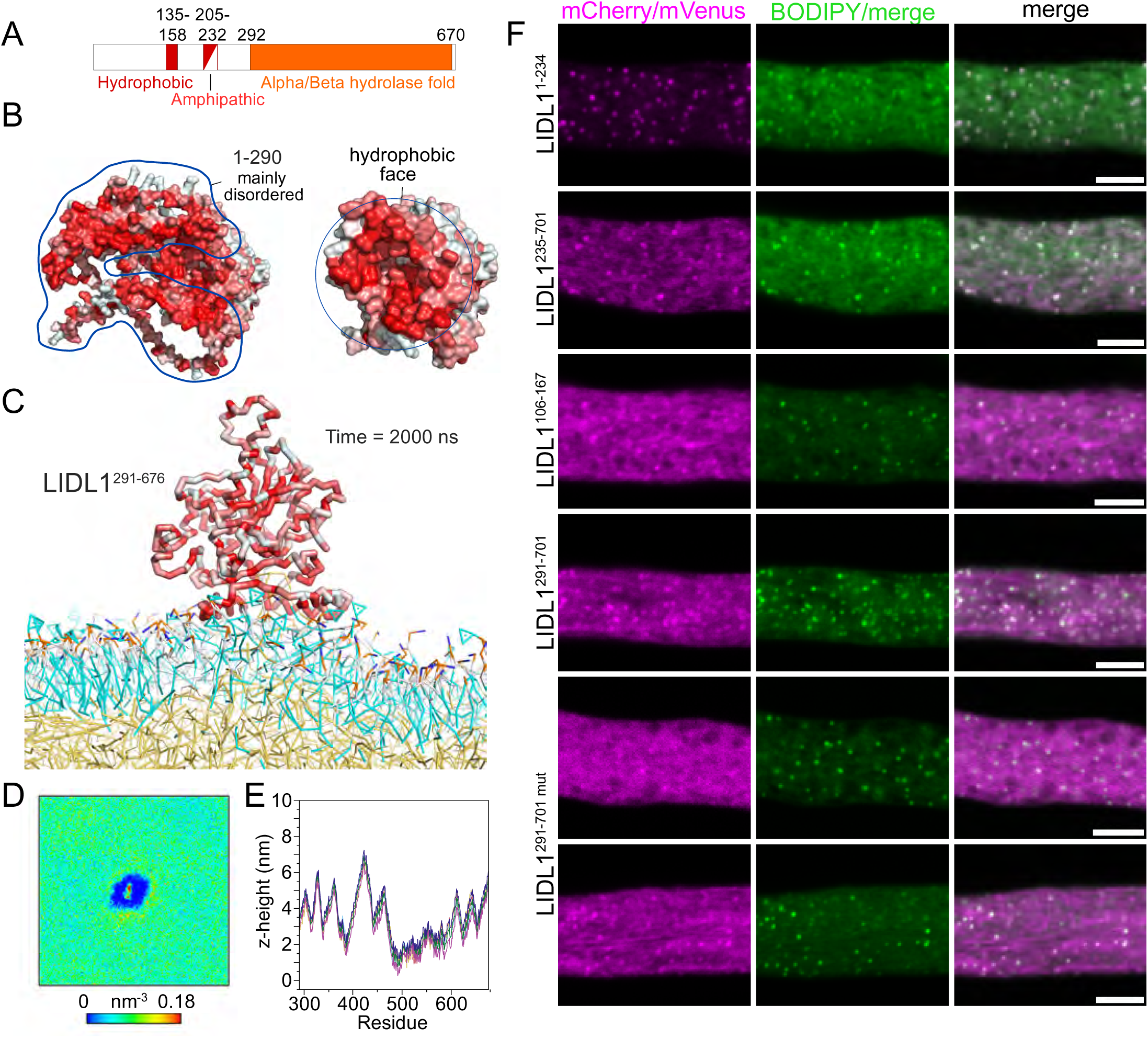
LIDL1 has two independent LD-targeting regions. **A** LIDL1 has a hydrophobic and an amphipathic helix at the N-terminus (res. 135-158 and 205-232) and an annotated Alpha/Beta hydrolase fold at the C-terminus (res. 292-670). **B** Surface representation of AlphaFold2 models of full-lenght LIDL1 (left panel) and LIDL1^291-676^ (right panel). The proteins are colored according to the Eisenberg hydrophobicity scale. Red, hydrophobic; white, hydrophilic. **C** Last frame of a representative MD simulation, in which LIDL1^291-676^ bound to the LD surface via the hydrophobic face within the simulation timeframe (time = 2000 ns). LIDL1^291-676^ was in contact with the LD system surface in 6 independent simulations. **D** Per residue z-height results of successfully bound simulations. Results from independent simulations are shown in different colors. **E** Phosphate density map of the binding leaflet in the last 100 ns of one representative simulation. **F** Subcellular localization of truncated and mutagenized LIDL1 proteins in tobacco pollen tubes. The fluorescent tag was attached to the C-terminus of LIDL1 variants. LDs were stained with MDH (monodansylpentane, for mVenus-tagged proteins) or BODIPY 493/503 (for mCherry-tagged proteins). Pollen tubes were incubated for 5-7 h before imaging. Scale bars, 5 µm.

Next, we generated a series of truncated and mutagenized LIDL1 proteins fused with mVenus or mCherry and transiently expressed them in tobacco pollen tubes. Consistent with the MD simulation result, LIDL1^235-701^ and LIDL1^291-701^ co-localized with LDs. Mutagenesis of 7 large hydrophobic amino acid residues to valine did not completely abolish the LD targeting of LIDL1^291-701^. However, in 5 out of 10 randomly selected transformed pollen tubes, the mutagenized LIDL1^291-701^ was mostly cytosolic (Figure 9 F), which was not observed in any pollen tubes transformed with non-mutagenized LIDL1^291-701^. These results suggest that the LD binding efficiency is affected by the large hydrophobic amino acid residues on the hydrophobic surface.

To probe the LD targeting capabilities of the N-terminus, which we could not simulate, we also tested two N-terminally truncated variants, LIDL^1-234^ and LIDL^106-167^. Interestingly, LIDL1^1-234^, and even the shorter hydrophobic helix, LIDL1^106-167^, also co-localized to the LDs in tobacco pollen tubes (Figure 9 E) indicating that LIDL1 has two independent targeting regions.

### Results from MD simulations can be an indicator for LD association *in vivo*

In this work, we mostly used MD simulations to investigate the binding mode of LD-associated proteins. Furthermore, the simulations predicted protein regions involved in binding that could be experimentally tested and verified. Next we wanted to test if this approach can also be used to pre-screen a number of LD-protein candidates as obtained for example by a proteomic approach (Scholz et al., 2026; Omata et al., 2024; Bouchnak et al., 2023; Dabisch et al., 2026; Chen et al., 2023). We performed such a screen retrospectively on a number of proteins that were found in the proteome of root LDs and were then cloned and tested by two cell biological approaches for LD association (Scholz et al., 2026).

From these proteins, fourteen different candidate LD proteins that covered a broad range of protein structures were selected and simulated for 2000 ns independent of protein size (Figure10 A). From our previous simulations we observed that LD binding proteins reached a high number of contacts in more than 5 out of 10 simulations and that the binding orientation was reproducible (Figures 1-6, 8, 9, S2-S4, S10), while this was not the case for the two control proteins tested (Figures S5 and S6). To score these parameters in an unbiased manner for the root LD proteins, we calculated two data points per protein. First, we calculated the number of contacts for each protein in all 10 simulations as an average of the last 50 ns, during which the number of contacts is expected to stabilize. We then took the average of only the five highest values to exclude simulations where the proteins were by chance not able to bind to the monolayer. Second, we calculated the pair-wise Pearson correlation *r* between all combinations of these five simulations in respect to their z-height plots and averaged the values. The result indicates how reproducible the protein orientation is during the interaction with the monolayer surface. With one exception, all root LD proteins had more than 350 contacts and a Pearson correlation of more than 0.9, close to the value of 1, which indicates that in all 5 simulations the proteins bound the LD surface in almost exactly the same orientation (Figure 10 B).

**Figure 10.**
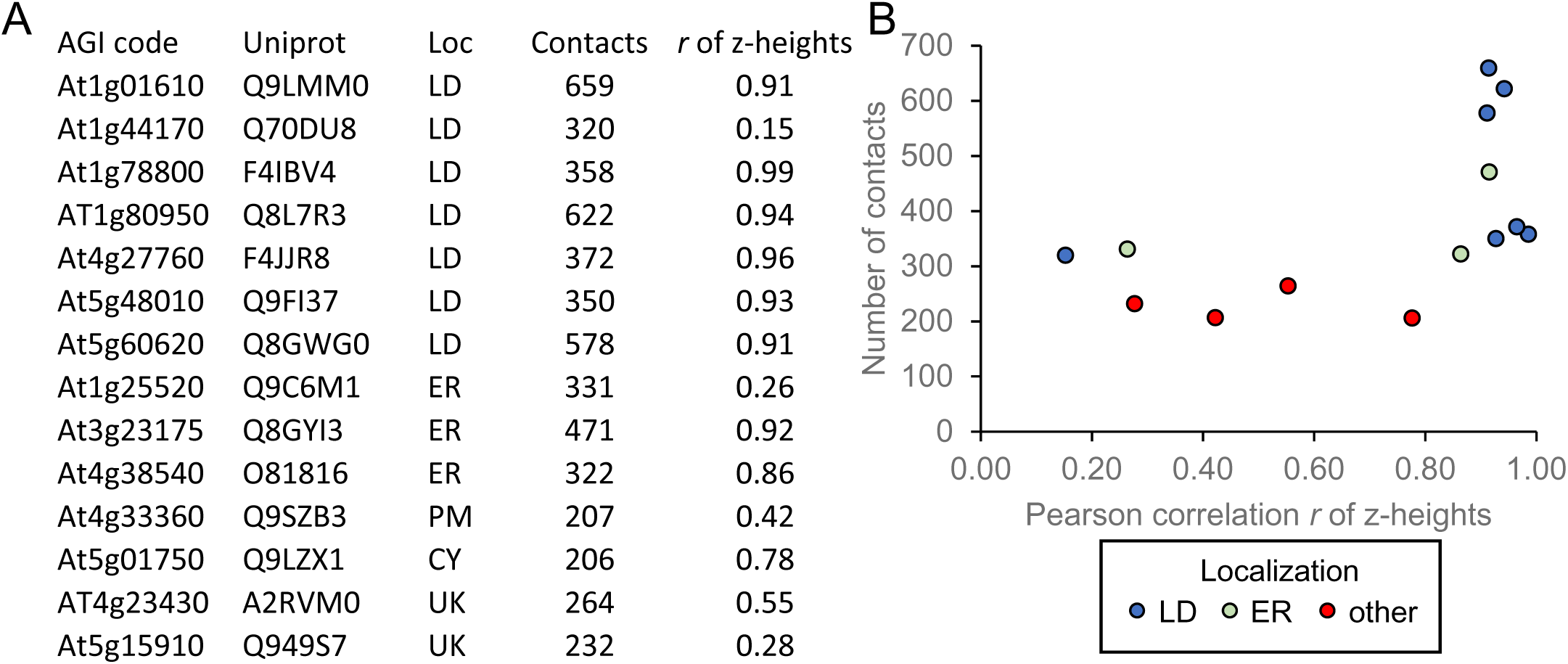
LD proteins have more contacts and bind in a more reproducible manner in MD simulations than non-LD proteins. **A** Selection of proteins found enriched in LD fractions from roots that were then tested in two cell biological systems for LD targeting (Scholz et al. 2026). Displayed are their localization (Loc) *in vivo* in Scholz et al. 2026 (PM, plasma membrane; CY, cytosol; UK unknown). Contacts and *r* of z-heights are derived from MD simulations of full-length versions of these proteins as follows: The average number of contacts during the last 50 ns of all simulations were calculated (see figures S2-S6 for examples). Only the five highest values from ten independent simulations were considered and averaged. The reproducibility of the binding orientation was estimated by calculating the Pearson correlation *r* between the z-heights of all combinations of these five simulations (See Figures 1-9 for examples) and taking the average of all *r* scores. **B** Graphical overview of the values of the tested proteins.

Two of three ER localized proteins had similar scores to the LD proteins. The other four proteins had much lower *r* scores and number of contacts, indicating that even with these simple metrics MD simulations can partially predict if a protein binds to LDs *in vivo*. When performed for a single target of interest, the other MD simulation results presented throughout the paper could give an even more detailed picture upfront.

All individual results of the simulations including the z-height and the number of contacts can be found in the Zenodo repository doi.org/10.5281/zenodo.15235178.

## Discussion

### OLE1, CLO1 and HSD1 may not target LDs via the historically proposed targeting mechanism

A proline knot/knob is a protein domain described as a hydrophobic hairpin with central conserved proline residues that inserts into the hydrophobic core of LDs to anchor proteins there (Chapman et al., 2012). Originally, based on the primary structure alone, oleosins and caleosins were proposed to have a proline knot in the middle of the protein flanked on both ends by other protein regions or domains (Huang and Huang, 2017; Jiang and Tzen, 2010; Frandsen et al., 2001). Steroleosins such as HSD1 were also hypothesized to harbor a similar but shorter hairpin termed proline knob at the N-terminus (Lin et al., 2002). However, the AlphaFold2 models and/or the targeting mechanism shown in MD simulations (Figures 2, 4, 5) suggest that not all proteins with a predicted proline knot or knob actually bind in the hypothesized orientation.

For OLE1 (Figure 2 A), the AlphaFold2 model remarkably resembles the structure proposed 35 years ago (Qu and Huang, 1990). The central hydrophobic hairpin with proline residues at the turning point can also be clearly observed in its AlphaFold2 model. This region alone was also able to target LDs in tobacco pollen tubes (Figure 2 F). However, in our MD simulations, this region embedded horizontally into the PL monolayer and not vertically reaching the neutral lipid core (Figure 2 B). It should be noted that the simulations did not include the flanking regions and placed OLE1 in the aqueous phase even though oleosins are known to bind to the ER first. These deviations could be the reason for the different binding orientation. However, the helix also contains twelve conserved hydrophilic serine and threonine residues that could make a horizontal binding energetically favorable. Furthermore, in contrast to transmembrane helices spanning a bilayer, a hairpin spanning a monolayer is not anchored at the other end of the membrane through strong interactions with the aqueous environment.

For CLO1 and HSD1, the AlphaFold2 models were clearly different from the previous proposed structures (Chapman et al., 2012). No obvious hairpin structure could be observed in the AlphaFold2 model of CLO1. In MD simulations, the annotated proline knot of CLO1 was in contact with the LD surface (Figure 5). However, instead of a hairpin anchor, it was part of a hydrophobic face. Mutagenesis of large hydrophobic residues within the hydrophobic surface led to an altered subcellular localization of CLO1 while previous mutations of the proline residues did not (Müller et al., 2017). For HSD1, the predicted hydrophobic hairpin domain, termed the proline knob (residues 1-40) was shown as a bent α-helix at the N-terminus in the AlphaFold2 model (Lin et al., 2002). In MD simulations, this region was also not inserted into the hydrophobic core but was embedded in the PL monolayer like an amphipathic helix (Figure 4). When being transiently expressed in tobacco pollen tubes, HSD1^1-46^ was both necessary and sufficient for LD targeting. We cannot exclude that there are limitations to the MD simulation system, which do not allow the helix to be inserted into the hydrophobic core of LDs, such as a limited sampling time which could prevent the system from overcoming a potential free energy barrier of insertion. Nevertheless, the MD simulations support the hypothesis that the binding orientation of CLO1 and HSD1 might be different from previous hypotheses.

### Proteins might target LDs via protein-protein interactions

Our MD simulation setup can only suggest a binding orientation of a protein that directly and strongly interacts with the LD surface. If such a direct and strong interaction is missing, the protein might still interact with the LD surface during the simulation, however, without establishing a reproducible binding orientation and with a lower number of contacts. As this was the case for LIME1 (Figure 7, Figure S11), we assume that it might not bind LDs *in vivo* in a self-sufficient manner but establish persistent LD localization via protein-protein interactions, as it presumably acts in a common pathway with other LD-localized proteins for the production of furan-ring-containing fatty acids with other LD-associated proteins (Omata et al., 2024).

### Proteins can target LDs via a hydrophobic face

Besides the previously described hydrophobic hairpin and amphipathic/hydrophobic helix, we suggest a third LD targeting structure: a hydrophobic surface of a folded domain which we termed the hydrophobic face.

Based on our simulations, this region of the protein inserts into the membrane just below the PL head groups of the LD surface, pushing these to the side (Figures 4-6 and 7-8). Its hydrophobic amino acid residues thereby do not interact with the LD core but the acyl chains of the PLs. On the LD-interacting hydrophobic face, more than 40 large hydrophobic amino acids can interact with LD PLs in case of OBL1 (Figure 6), however, less are required for the initial binding initiation indicated by our mutagenesis of twelve large hydrophobic residues within region 436-495 that changed OBL1’s LD localization. Such amino acids were shown to be crucial on several amphipathic helices and a variety of their muteins for detecting and inserting into membrane packing defects on the LD surface (Olzmann and Carvalho, 2019; Prévost et al., 2018). Contrary to these helices which were proposed to only fold while inserting into packing defects, we expect the hydrophobic faces to be fully folded before reaching the LD surface. Therefore, we assume that their mutagenesis to smaller hydrophobic amino acids (valine or alanine), which led to a depletion of LD binding of these proteins in tobacco pollen tubes, altered the subcellular localizations by a reduced ability to detect packing defects and not by a changed fold or the inability to fold once in contact with the LD surface.

The hydrophobic face may not only be a structural feature for LD targeting but could have a decisive advantage over other binding modes: it might give enzymes access to hydrophobic substrates inside the LDs, thus playing a role in catalysis of hydrophobic substrates stored in LDs. For example, CAS1 has a homolog in the human genome, lanosterol synthase, also referred to as oxidosqualene cyclase (HsOSC, Uniprot code P48449). AtCAS1 shares 41.7% identity and its AlphaFold2 model shows strong structural similarity with HsOSC. HsOSC has a channel for substrate passage in the center of the HsOSC membrane binding region (Thoma et al., 2004). In Arabidopsis, CAS1 catalyzes the synthesis of cycloartenol from oxidosqualene (Corey et al., 1993) and a substrate channel similar to the one of HsOSC can be observed that originates in the middle of the hydrophobic face (Figure S13). Thus, it is possible that AtCAS1 is taking up its substrate through this hole in the hydrophobic face. Interestingly, also other proteins with hydrophobic faces, such as CLO1, UFAO1, OBL1, LIDL1 and LDAH1, are reported or are annotated to have catalytic activity (Müller and Ischebeck, 2018; Poxleitner et al., 2006; Omata et al., 2024; Mistry et al., 2021), while it is not found in presumably structural or scaffold proteins like oleosins, SLDPs, PUX10 or MYOB14. In most MD simulations, in which UFAO1, OBL1, LIDL1 and LDAH1 reached the LD surface, typically a PL or occasionally also a TAG molecule entered the potential active side of these proteins (see Figures 4 J, 6 A, 9 C and 8 J, respectively) leading to an increased PL density in the center of the binding region (see Figures 4 K, 6 B, 9 E and 8 K, respectively).

### Several LD targeting mechanisms can cooperate

Compared to their full-length counterparts, HSD1^1-47^, UFAO11-50 and LDAH1^1-50^ showed a more pronounced ER localization, suggesting that additional features contributing to LD targeting may exist (Figure S9). HSD1, UFAO1 and LDAH1 all have a hydrophobic face on their C-terminal domain in addition to their N-terminal hydrophobic helices (Figure 4, 8). Even though the C-terminal region of HSD1 and UFAO1 did not target LDs on their own *in vivo*, the hydrophobic face interacted with the LD surface in MD simulations. Thus, there is a possibility that these two proteins and LDAH1 first bind to the ER via their C-terminus while the N-terminus stabilizes LD localization after the protein has moved there.

### The LD targeting classification could be restructured and expanded

For proteins that target LDs via their structural features, two important pathways, class I and class II, have been described (Kory et al., 2016; Bersuker and Olzmann, 2017). Class I proteins either refers to proteins targeting to LDs via a hydrophobic hairpin (Bersuker and Olzmann, 2017), or proteins that are initially inserted into the ER membrane before being transferred to the surface of growing LDs that are still attached to the ER surface (Kory et al., 2016). Class II proteins are defined as proteins that target LDs via amphipathic helices (Bersuker and Olzmann, 2017) directly from the cytosol (Kory et al., 2016).

To properly include the hydrophobic face, which we identified as an LD-targeting structural feature, into the current classification system, we propose a more nuanced classification. Currently, the two classes are often defined both by the protein’s route to the LD and its structural feature that mediates the binding (Bersuker and Olzmann, 2017; Kory et al., 2016). While these are often linked and typically amphipathic/hydrophobic helices are thought to target LDs directly from the cytosol, our results for HSD1 and UFAO imply a slightly more complicated picture as discussed above.

To avoid inferring a route based on the structural feature alone, we propose a two layered classification system: As the first layer, class I and class II solely indicate the targeting route, with class I proteins first binding to the ER and class II proteins bind to LDs directly from the cytosol. As the second layer, we suggest distinguishing between the structural features that mediate the binding. These include hydrophobic hairpins or proline knots, amphipathic/hydrophobic helices, lipid anchors, protein-protein interactions and hydrophobic faces. This two-layered classification system allows for a more nuanced distinction between different LD-targeting proteins and acknowledges that the structural features of LD targeting might be well predicted or known while the exact route of binding remains to be elucidated.

### The use of MD simulations in studying LD function

Being originally considered to be inert storage depots, the number of functions associated with plant LDs has strongly increased over the years (Bouchnak et al., 2023; Guzha et al., 2023). One driving factor has been the identification of a large number of LD-associated proteins that added functions in regard to protein turnover, metabolism and LD positioning. In comparison to other organelles, no clear targeting signals have been identified for LD proteins, and most LD-targeting proteins were identified from proteomic screens combined with assaying individual proteins via cell biological tools.

Here, MD simulations are shown to be a valuable tool that could help to prescreen longer lists of candidates. Simulating several proteins like in this study required a total of around 700 µs of coarse-grained MD simulations with weeks of computing on a high-performance computing cluster. With an increase in available computing power and the trivially parallelable approach used here, however, it can be envisioned that hundreds of proteins could be screened based on genetic information alone to identify previously unknown LD-association of proteins that can then be confirmed through cell biology. Furthermore, with the help of machine learning algorithms, targeting probability based on primary or tertiary structure could aid in compiling a list of likely LD protein candidates. MD simulations can also help to understand how proteins interact with the LD surface including enzymes that obtain their substrates from the LD core or surface. This could be an advantage for enzyme design in synthetic biology when LDs are used as microfactories in biotechnology. In conclusion, MD simulations can greatly increase our understanding of LD biology when supported with *in vivo* experiments.

## Acknowledgments

We thank Ivo Feussner (University of Göttingen) for hosting and supporting the Ischebeck lab during parts of this work, and Jan-Ole Niemeier and Markus Schwarzländer for support with microscopy.

## Author contributions

K.C., S.T., M.B., and T.I. designed the research; S.S., P.N., P.W., and M.B. performed research; S.S., P.W., K.C., S.T. and M.B. analyzed data; S.S., P.W., M.B. and T.I. wrote the paper with the help of all authors.

## Supplemental Data

**Supplemental Figure S1.** Subcellular localization of AtPUX10^94-143^-mCherry in tobacco pollen tubes

**Supplemental Figure S2.** Closest distance and number of contacts of PUX10^94-143^.

**Supplemental Figure S3** Closest distance and number of contacts of SLDP1.3^19-81^.

**Supplemental Figure S4** Closest distance and number of contacts of OLE1.1^33-124^.

**Supplemental Figure S5** Closest distance, number of contacts and z-height of Lysozyme.

**Supplemental Figure S6.** Closest distance, number of contacts and z-height of CINV1.

**Supplemental Figure S7.** MD simulation of the N-terminus of MYOB14.

**Supplemental Figure S8.** Predicted structure of LDDH1.

**Supplemental Figure S9.** Several proteins and/or their truncated/mutated version showed co-localization with both the ER and LDs.

**Supplemental Figure S10.** Number of contacts of CLO1, CAS1 and OBL1.

**Supplemental Figure S11.** Number of contacts of LIME1.

**Supplemental Figure S12.** Subcellular localization of LDAH1.2 in tobacco pollen tubes.

**Supplemental Figure S13.** Cross-section of the predicted structure of CAS1 displaying the active site

**Supplemental Table S1** Overview of truncated and mutagenized protein variants.

**Supplemental Data S1** Glycerophospholipid composition of isolated LDs from Arabidopsis.

**Supplemental Data S2** Triacylglycerol composition of isolated LDs from Arabidopsis.

**Supplemental Data S3** Alignment of SLDP orthologs.

**Supplemental Data S4** Alignment of Oleosin orthologs.

## Data availability

All simulation data is available under the Zenodo repository doi.org/10.5281/zenodo.15235178.

