## Supplemental Table S1 for "Molecular dynamics simulations and *in vivo* binding studies indicate different binding modes of plant lipid droplet proteins including binding via a hydrophobic face"

**Supplemental table S1. The LD targeting sequence analyzed in this study.** The mutagenized amino acids on hydrophobic surfaces are highlighted in red.

| AGI No. | Abbreviation | Region | LD Targeting sequence |
|---|---|---|---|
| AT4G25140.1 | OLE1 | 33-124 | GSLLVLSSLTLVGTVIALTVATPLLVIFSPILVPALITVALLITGFLSSGGFGIAAITVFSWIYKYATG |
| AT1G65090.3 | SLDP1 | 19-81 | SWIWSKAVSVGKKVLTAGVVVSSAPLLVPSLFVASTLAFLSSVPFCLFLANYACTQKVMSTLL |
| AT4G10790.1 | PUX10 | 94-143 | NRGPGVAWRIITLPISIVSGSLGLVSGAIGLGIWAAGGVLSYSLGMLGFR |
| AT4G13160.1 | MYOB14 | 1-50 | MELLFLLCSIIYTSITSIFFSLLLPFRFLLHRLLPSRAAVDPNVSFYEGT |
| AT5G50600.1 | HSD1 | 1-50 | MELINDFLNLTAPFFTFFGLCFFLPPFYFFKFLQSIFSTIFSENLYGKVV |
| AT1G10740.1 | LDAH1 | 1-50 | MIKLWMMTSLQLAELFVSSIVHLLYGFYIFSSAVAGDISQTLNDYLFKSN |
| | | 228-281 | CLLQLCRIGVIMYDWLDISWLKTYYNFGFDHFNISWKKTGVRGLVDCLMGNAGPF |
| AT1G30130.1 | UFAO1 | 1-50 | MDYQESYRLTFYGILVAFMELAFAYCLLCVSAFVFITSKLLLFMPCIGYQ |
| AT1G75180.1 | LDDH1 | 240-315 | TEYGSYKQEYEGEHSDMYWNREYADGEDGEMSWFNAGVRVGVGIGLGVCVGLGIGVGLLVRT YQSTTRNFRRRII |
| AT3G14360.1 | OBL1 | 67-118 | DHRWVIVVSILVRKIIRLLRTPMEFTGFVVDFFLNLFSANGGFFGLLLRLIQ |
| | | 436-495 | EDEPDPNPYGLRYKILGHVIAVWELVRGLTMGYTHGPDYKEGWFRILFRLMGLVIPGLSD |
| AT4G26740.1 | CLO1 | 97-131 | NIIGSLIIAAVINLTLSYATLPGWLPSPFFPIYIH |
| | | 189-213 | IFGWIAGKIEWGLLYLLARDEEGFL |

| AGI No. | Abbreviation | Region | LD Targeting sequence |
|---|---|---|---|
| AT2G07050.1 | CAS1 | 239-251 | EIWLLPYFLPIHP |
| | | 315-336 | PLVQDILWASLHKIVEPVLMRW |
| | | 539-546 | PWLELINP |
| AT4G33110.1 | LIME1 | 1-45 | MEKIIDVAYGASVKVGLTLLEMNLLPDIVIRRLTRLLLAGRLRSG |
| | | 300-355 | DKEIVAVKKIMEVTYGKEEAVKWMVYWRTFFIAVAELFGYNNGEEWMVAHFLFKKK |
| AT1G18460.1 | LIDL1 | 479-539 | NMCFTLMEYTFLFLGPVLSRIVPAFYIPTKFFRMLLNKLARDFHNYPAVGGLVQTLMSYVV |
