## Supplemental Figures S1-S13 for "Molecular dynamics simulations and *in vivo* binding studies indicate different binding modes of plant lipid droplet proteins including binding via a hydrophobic face"

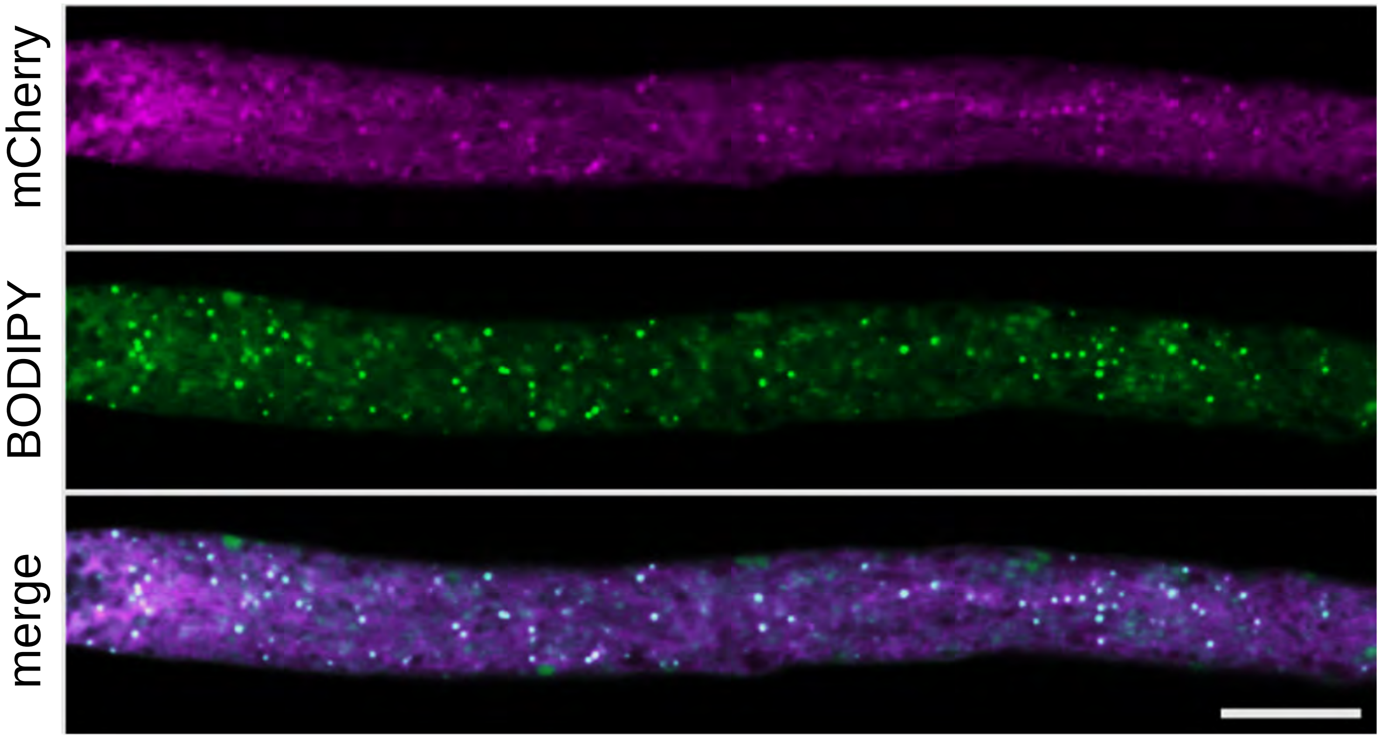


**Supplemental Figure S1. Subcellular localization of AtPUX10$^{94\text{-}143}$-mCherry in tobacco pollen tubes.**

AtPUX10$^{94\text{-}143}$ with C-terminally tagged mVenus was transiently expressed in tobacco pollen tubes. The pollen tubes were cultivated for 5-7 h and fixed with formaldehyde before imaging. LDs were stained with BODIPY 493/503. Scale bar, 10 nm.

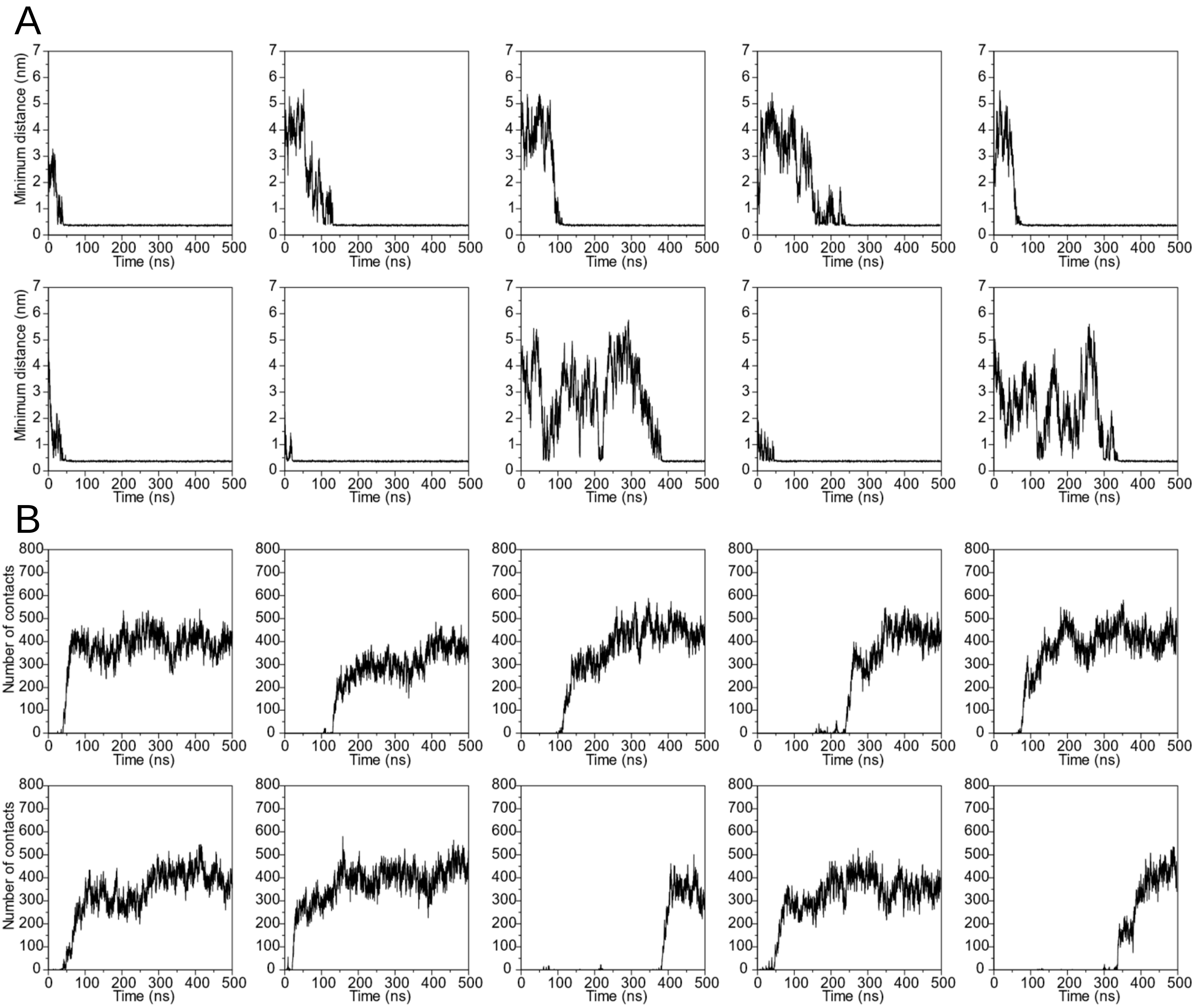


**Supplemental Figure S2 PUX10$^{94\text{-}143}$ interacts with the LD system with a high number of contacts in coarse-grained MD simulation.**

Data derives from ten individual simulations. **A** The closest distance of the protein to the LD surface, and **B** the number of contacts (the number of protein-LD bead pairs that are closer than 0.6 nm to one another) was mapped over time.

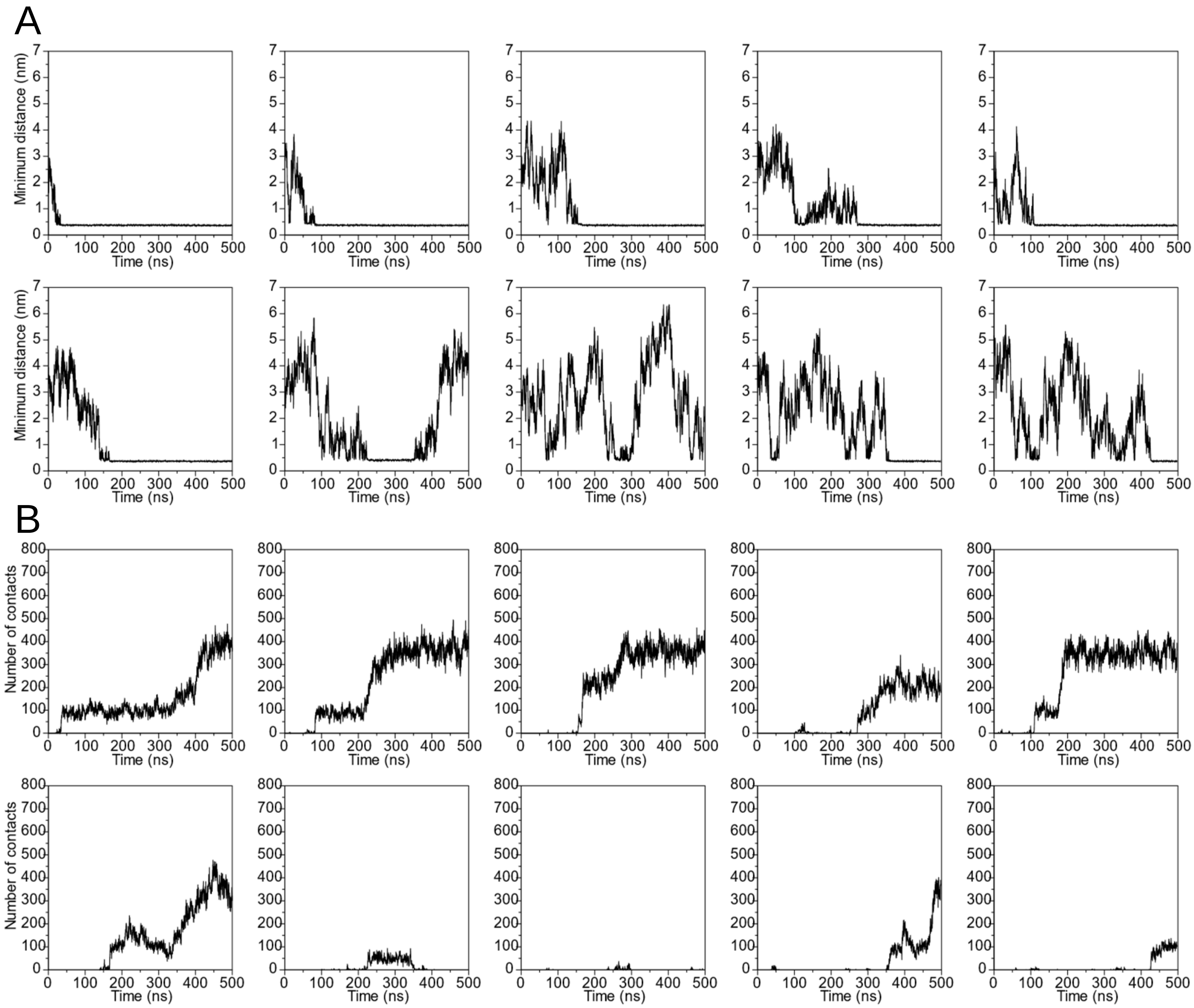


**Supplemental Figure S3 SLDP1.3$^{19-81}$ interacts with the LD system with a high number of contacts in coarse-grained MD simulation.**

Data derives from ten individual simulations. **A** The closest distance of the protein to the LD surface, and **B** the number of contacts (the number of protein-LD bead pairs that are closer than 0.6 nm to one another) was mapped over time.

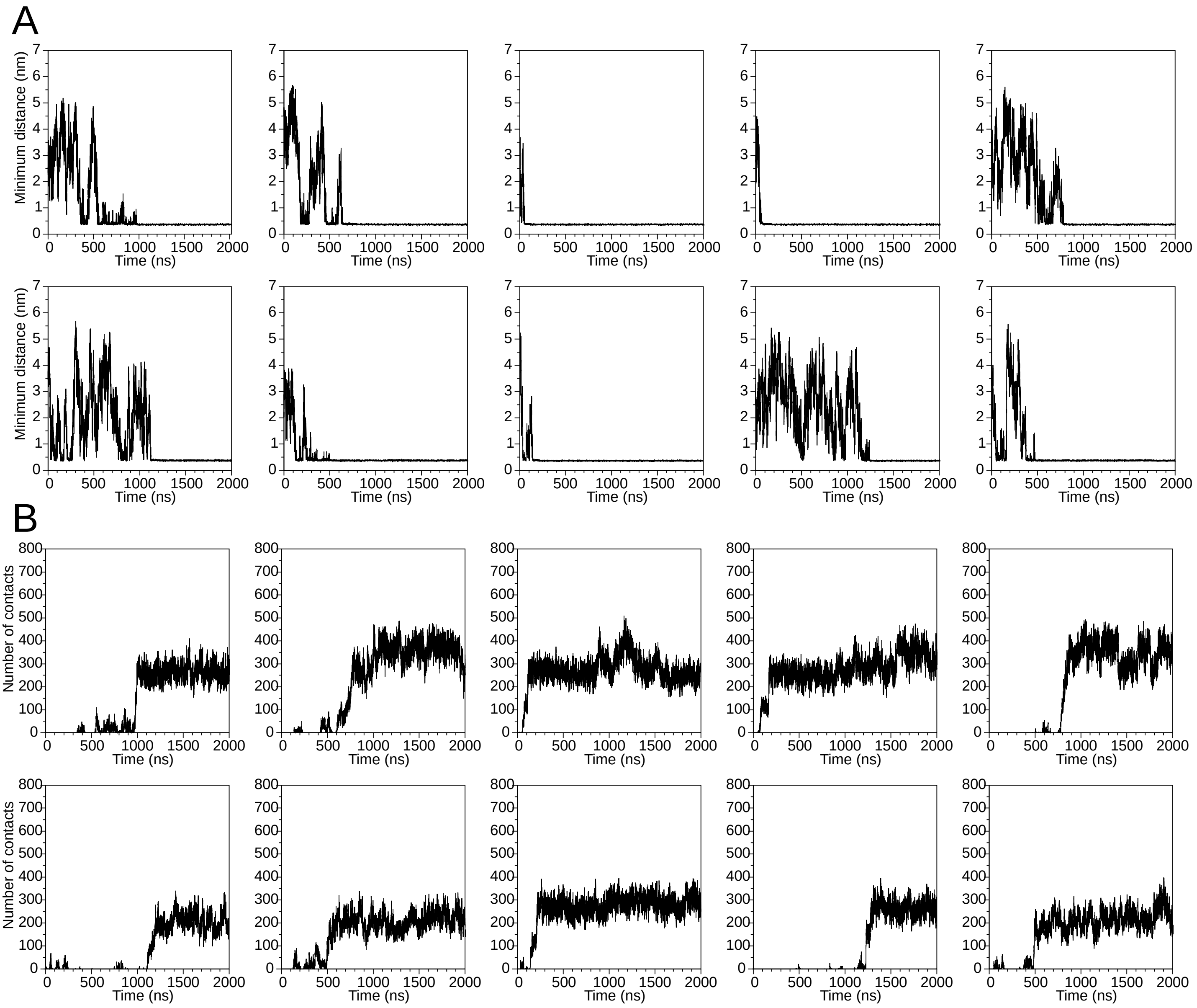


**Supplemental Figure S4 OLE1.1$^{33\text{-}124}$ interacts with the LD system with a high number of residues in coarse-grained MD simulation.**

Data derives from ten individual simulations. **A** The closest distance of the protein to the LD surface, and **B** the number of contacts (the number of protein-LD bead pairs that are closer than 0.6 nm to one another) was mapped over time.

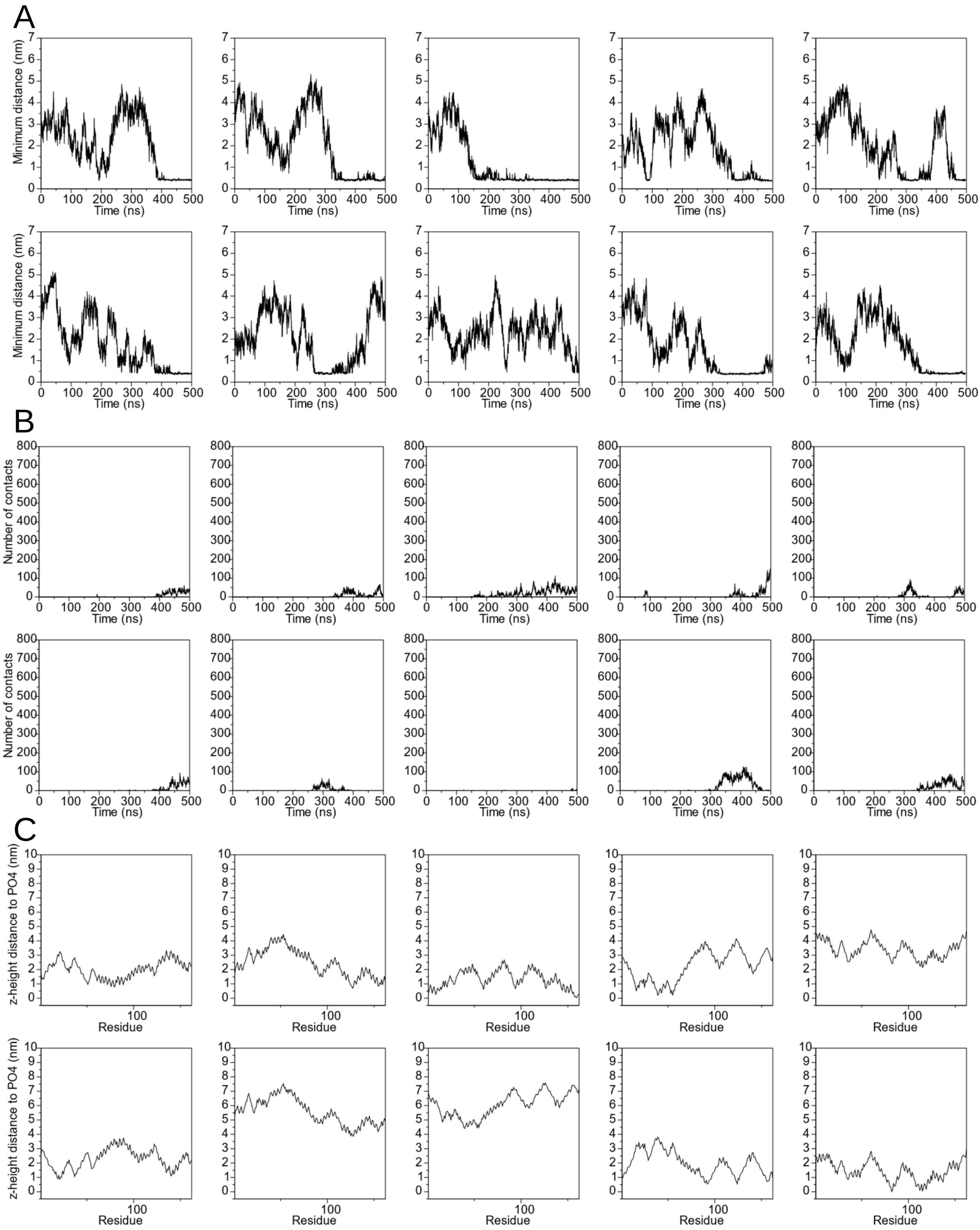


**Supplemental Figure S5 Lysozyme interacts with the LD surface only via few contacts and in random orientation.**

Data derives from ten individual simulations of the enzyme lysozyme from *Enterobacteria phage* T4. **A** The closest distance of the protein to the LD surface, and **B** the number of contacts (the number of protein-LD bead pairs that are closer than 0.6 nm to one another) was mapped over time. The number of contacts was low in all cases and binding sometimes transient. **C** Per residue z-height distance to the averaged phosphate z-height of the closest LD system leaflet at the end of the simulations.

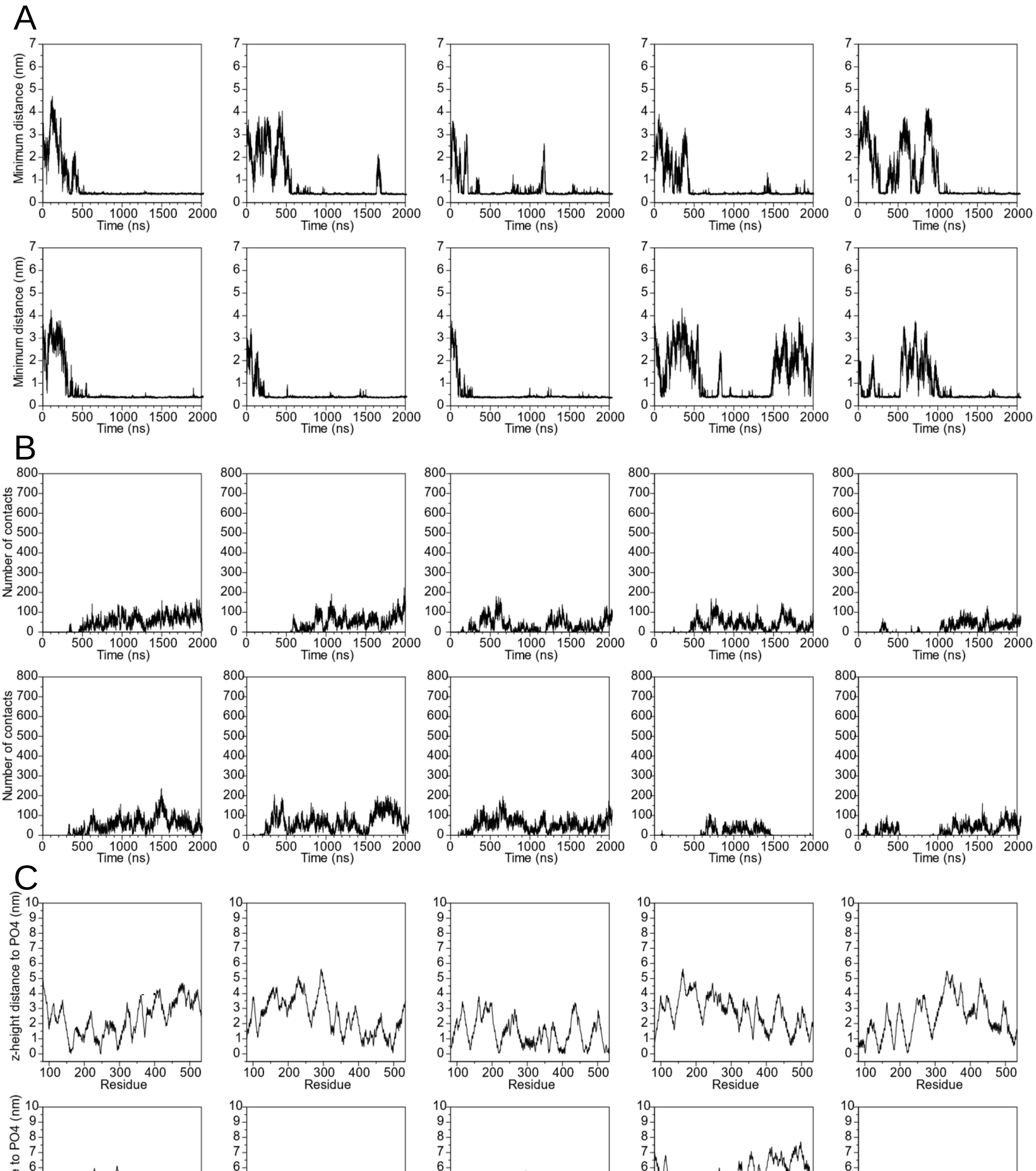


**Supplemental Figure S6 CINV1 interacts with the LD surface only via few contacts and in random orientation.**

Data derives from ten individual simulations. **A** The closest distance of the protein to the LD surface, and **B** the number of contacts (the number of protein-LD bead pairs that are closer than 0.6 nm to one another) was mapped over time. The number of contacts was low in all cases and binding sometimes transient. **C** Per residue z-height distance to the averaged phosphate z-height of the closest LD system leaflet at the end of the simulations.

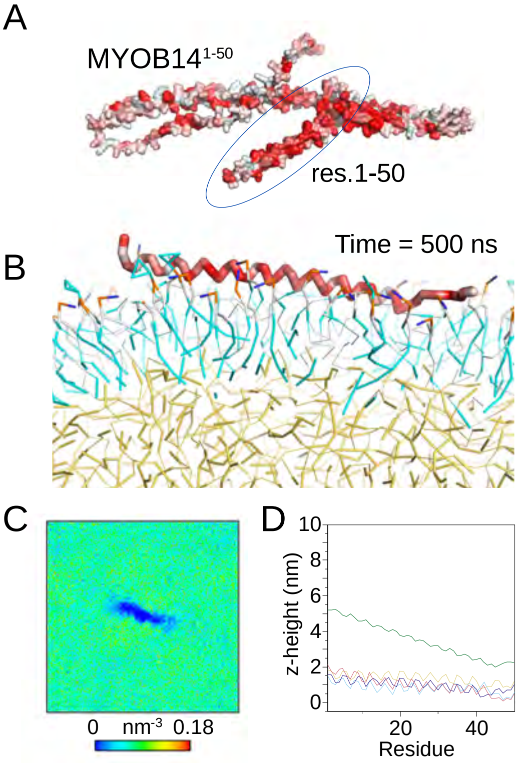


**Supplemental Figure S7. The N-terminus of MYOB14 binds to LDs in MD simulations.**

**A** AlphaFold2 model of MYOB14 displayed in surface representation colored according to the Eisenberg hydrophobicity scale. Red, hydrophobic; white, hydrophilic. MYOB14$^{1\text{-}50}$ arrived at the LD system in 6 independent simulations during the simulation timeframe (time = 500ns). **B** Last frame of a representative simulation, in which MYOB14$^{1\text{-}50}$ bound to the LD system. **C** Averaged phosphate density of the binding leaflet in the last 100 ns of one representative simulation. MYOB14$^{1\text{-}50}$ pushed the PL head groups at the protein binding site away, suggesting that it was embedded into the PL monolayer in the simulations. **D** Per residue z-height distance of the 6 successfully bound simulations. Results from independent simulations are shown in different colors.

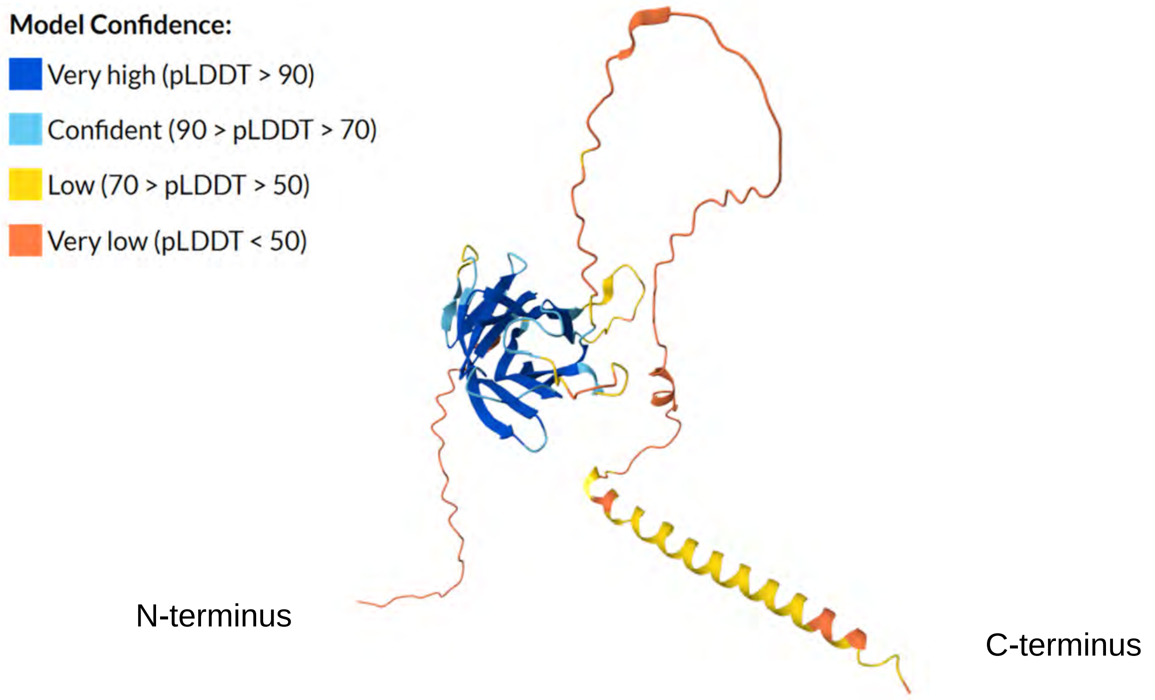


**Supplemental Figure S8. The full-length version of LDDH1 contains long unordered regions which makes full length simulation difficult.**

**A** AlphaFold2 model of LDDH1 displayed in cartoon representation colored according to the model confidence score. According to the model, the residues 270-315 form an α-helix. This helix alone can bind to LDs both *in silico* and *in vivo*.

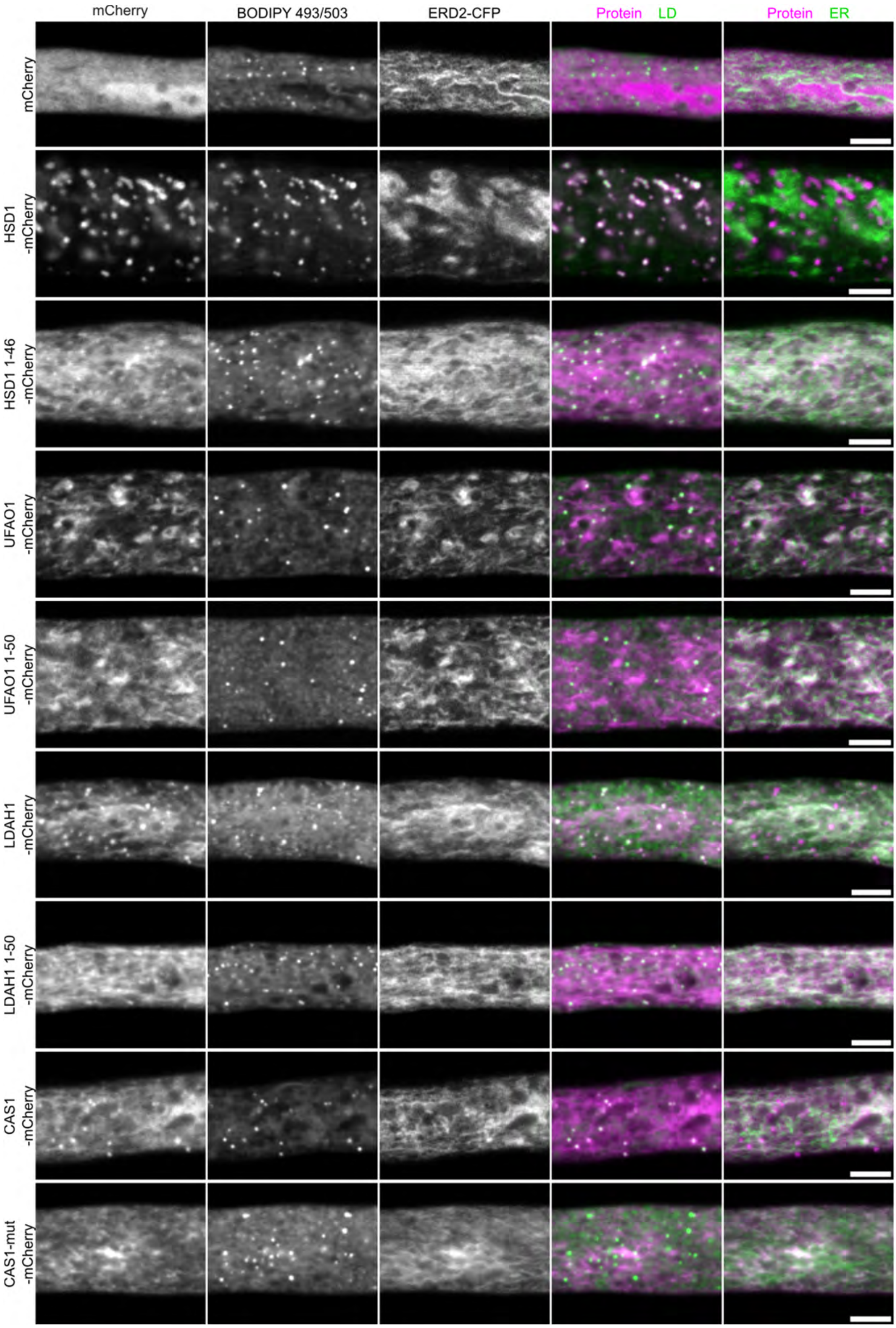


**Supplemental Figure S9. Several proteins and/or their truncation/mutated protein showed co-localization with both the ER and LDs.**

Proteins/truncations with observed network-like structures were co-expressed with the ER marker, ERD2-CFP in tobacco pollen tubes. Pollen tubes were incubated for 5-7 h before imaging. An image representative of at least 10 pollen tubes is shown in each row. LDs were stained with BODIPY 493/503. Scale bar = 5nm.

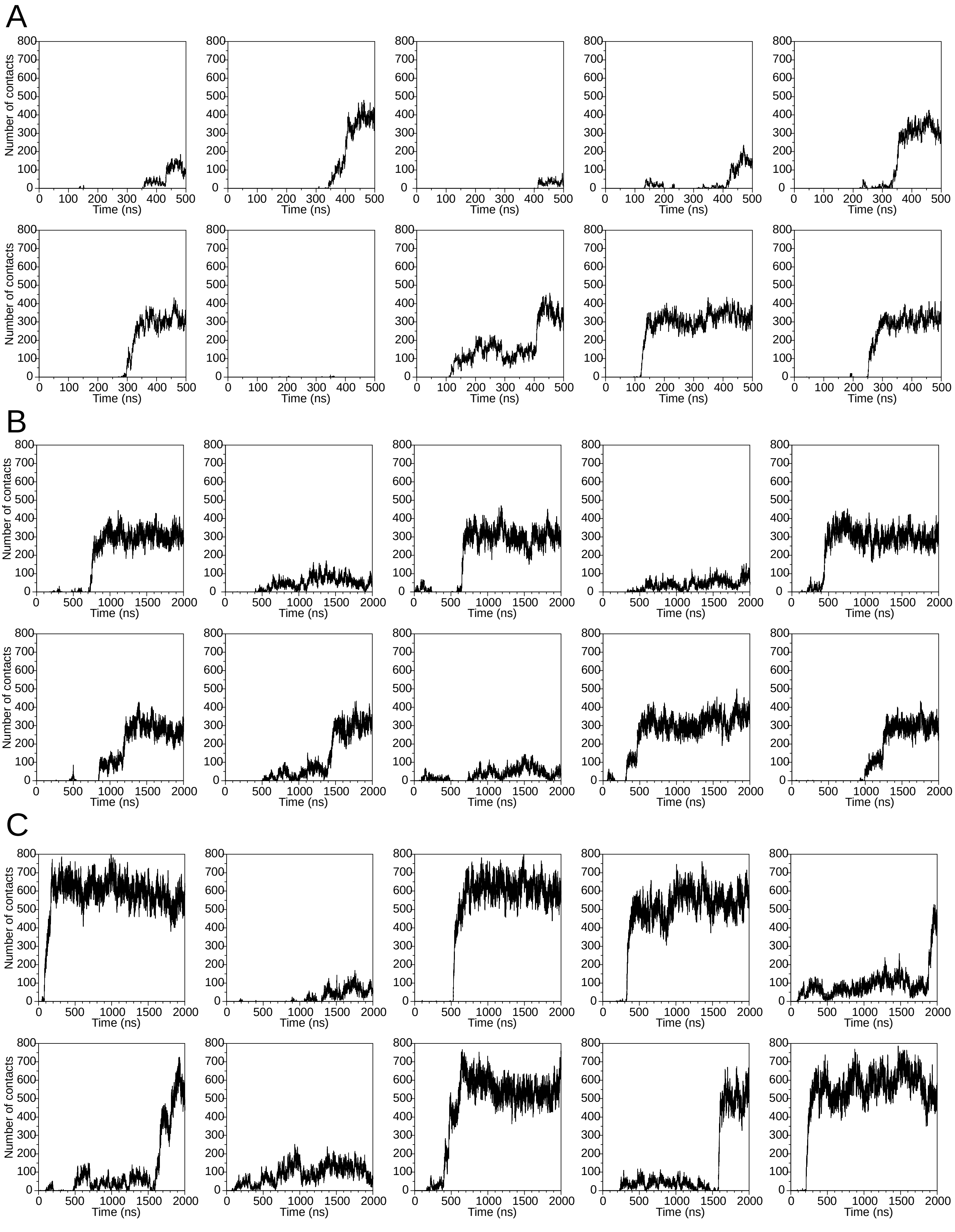


**Supplemental Figure S10 Proteins can bind the LD surface through a hydrophobic face with a high number of contacts.**

The number of contacts (the number of protein-LD bead pairs that are closer than 0.6 nm to one another) was mapped over time for the proteins **A** CALEOSIN1 (CLO1), **B** CYCLOARTENOL SYNTHASE 1 (CAS1) and **C** OIL BODY LIPASE 1 (OBL1). In case of binding the number of contacts was stable, high and comparable between simulations of the same proteins.

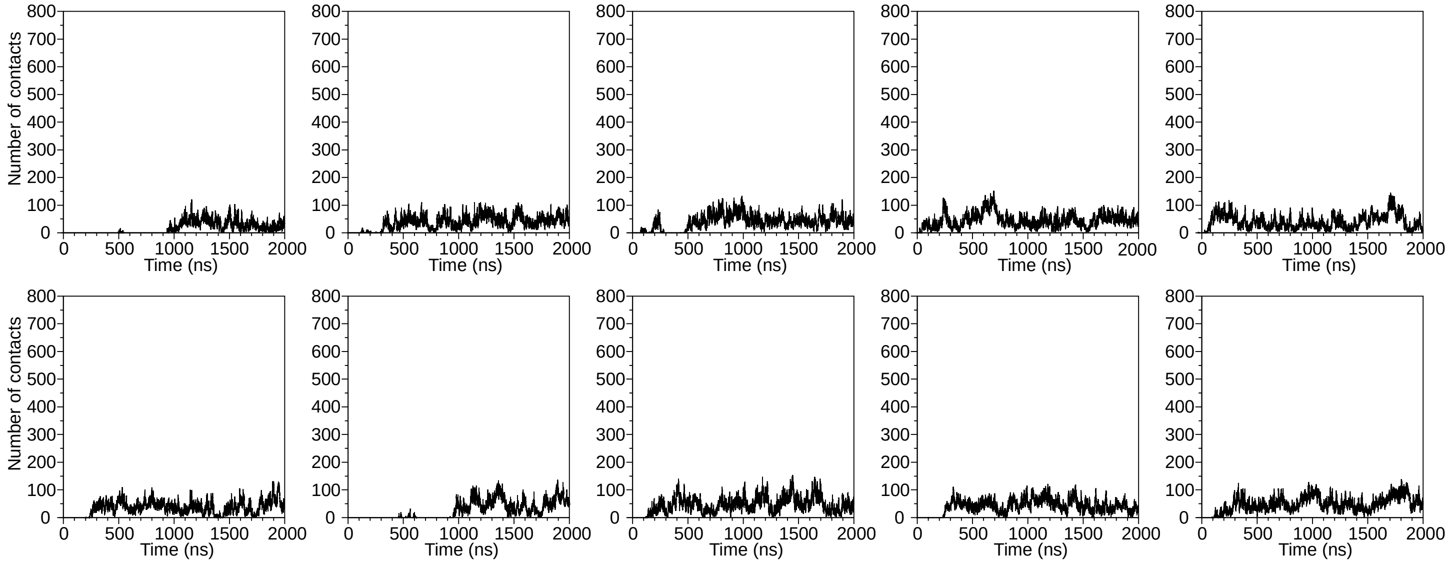


**Supplemental Figure S11 LIME1 interacts with the LD surface only via few interactions.**

Data derives from ten individual simulations. The number of contacts (the number of protein-LD bead pairs that are closer than 0.6 nm to one another) was mapped over time. The number of contacts was low in all cases.

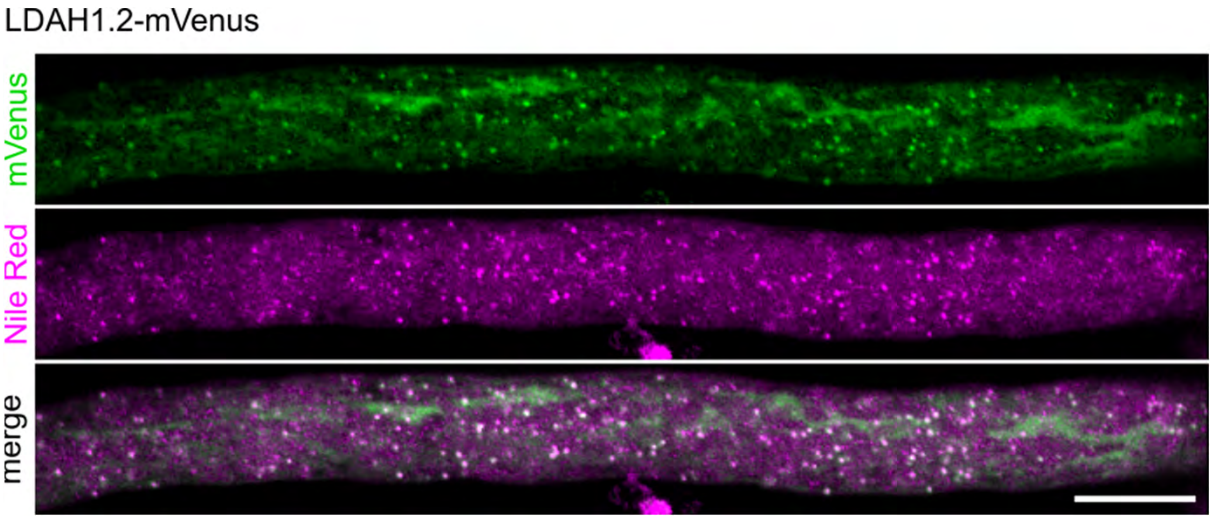


**Supplemental Figure S12 Subcellular localization of LDAH1.2 in tobacco pollen tubes.**

LDAH1.2 (missing res. 406-473 of LDAH1.1) with C-terminally tagged mVenus were transiently expressed in tobacco pollen tubes. The pollen tubes were incubated for 5-7 hours and fixed with formaldehyde before imaging. LDs were stained with Nile Red. Scale bar = 10 nm.

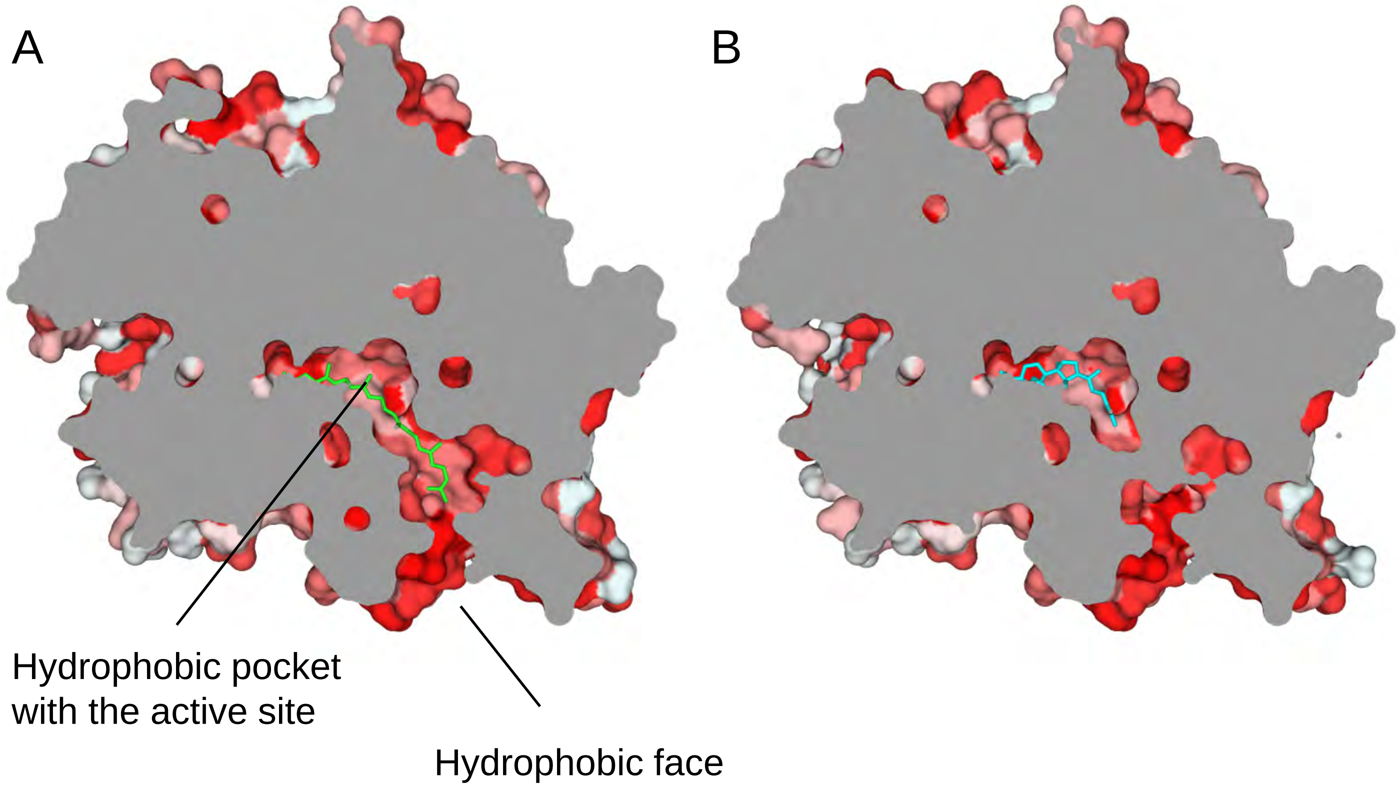


**Supplemental Figure S13 The hole in the hydrophobic face of CAS1 leads to the active site**

Cross section of CAS1 structures predicted with AlphaFold3 together with **A** the substrate (green) or **B** the product (light blue). The protein is colored according to the Eisenberg hydrophobicity scale. Red, hydrophobic; white, hydrophilic.
